# Ultra-secure storage and analysis of genetic data for the advancement of precision medicine

**DOI:** 10.1101/2024.04.16.589793

**Authors:** Jacob Blindenbach, Jiayi Kang, Seungwan Hong, Caline Karam, Thomas Lehner, Gamze Gürsoy

## Abstract

Cloud computing provides the opportunity to store the ever-growing genotype-phenotype data sets needed to achieve the full potential of precision medicine. However, due to the sensitive nature of this data and the patchwork of data privacy laws across states and countries, additional security protections are proving necessary to ensure data privacy and security. Here we present SQUiD, a **s**ecure **qu**eryable **d**atabase for storing and analyzing genotype-phenotype data. With SQUiD, genotype-phenotype data can be stored in a low-security, low-cost public cloud in the encrypted form, which researchers can securely query without the public cloud ever being able to decrypt the data. We demonstrate the usability of SQUiD by replicating various commonly used calculations such as polygenic risk scores, cohort creation for GWAS, MAF filtering, and patient similarity analysis both on synthetic and UK Biobank data. Our work represents a new and scalable platform enabling the realization of precision medicine without security and privacy concerns.

## 2 Introduction

Precision medicine aims to tailor medical care to the characteristics of an individual’s unique genetic makeup, lifestyle, and environment. This approach has garnered considerable attention worldwide due to its potential to enhance patient outcomes and mitigate healthcare expenses[33]. But, several significant obstacles impede the realization of the full potential of precision medicine. One such challenge is the need for extensive and diverse patient genotype-phenotype datasets in order to advance the diagnosis and treatment of future patients [69]. However, this need for large amounts of data is often in conflict with the need to protect patient privacy [6]. This challenge is further complicated by the heterogeneous regulatory landscape governing privacy protection, with varying definitions and practices across different jurisdictions (*e.g*., General Data Protection Regulation [GDPR] in Europe *vs*. frameworks in USA) [27,67]. Furthermore, individual hospitals and institutions maintain their own policies due to the prevalence of health data breaches and privacy attacks.

Genomic data plays a pivotal role in precision medicine research, enabling the customization of medical care based on specific genetic variants, biomarkers, and inherited traits. Thus, there is a surge in data generation, which has challenged the ability of local servers to accommodate the rapid growth of data size and increased computational requirements [66]. Therefore, there is a pressing need and significant push towards cloud computing. However, this exacerbates the concerns about the privacy and prohibitions on use of personal data due to local, global, and/or institutional privacy policies. For example, with the introduction of the GDPR in Europe, the storage of genomic and related data in the cloud has become more stringent with the requirement of appropriate security measures in place such as encryption. Starting from 2023, many states in the US (California, Connecticut, Colorado, Utah, and Virginia) are entering a new GDPR-like privacy era that will have similar requirements about storing genomic and related data in the cloud [4]. Yet, the current state of privacy preservation through laws and institutional policies is fraught with instability and unpredictability, which poses significant challenges to the research community. If the data is kept in the encrypted form in cloud servers, then researchers, who are approved for access, need to download large quantities of data locally and decrypt them to perform analysis, which defeats the purpose of outsourcing the storage to the cloud. This situation creates additional hurdles for scientists, especially when attempting to combine multiple data sets to gain statistical power. Furthermore, it creates significant delays in research and requires large amounts of resources, which impedes the democratization of data access. As a result, advances in medicine will significantly be impacted if new privacy-preserving frameworks that comply with laws and policies are not developed and implemented.

Homomorphic encryption is one of the cryptographic tools that enables direct computations of functions on encrypted data in the public cloud. Homomorphic encryption has emerged as a useful approach to keep the data secure at rest, at transit, and during analysis. But, this approach also has thus far presented severe bottlenecks in its applicability, scalability, and performance [50,2]. However, recent advances in algorithm designs and computing power have enabled an increase in the use of homomorphic encryption in genomics. For example, it has been shown that privacy-enhancing genome-wide association studies (GWAS) can be possible [62,70,13,44]. It has also been shown that secure genotype imputation is feasible and scalable using homomorphic encryption [71,20,36]. Homomorphic encryption was also used for genomic variant querying [19], regression analysis for rare disease variants [68], and inference using genetic variants in machine learning applications [60]. These methods have added tremendous algorithmic advances to the field and paved the way for more practical privacy-preserving analysis of genomes. However, their use in genotype-phenotype database settings has been limited. This is primarily attributed to two factors. Firstly, homomorphic encryption relies on public key cryptography, which is designed for client-server scenarios where the client owns the dataset and delegates computation to the cloud. However, in the context of genotype-phenotype databases, the data owner encrypts the data while multiple researchers access and analyze it. Secondly, the computational burden associated with homomorphic encryption makes it infeasible for applications involving large sample sizes. Both the storage size of encrypted data (*i.e*., ciphertexts) and the computation times for homomorphic encryption are several orders of magnitude greater than those for the original plaintexts [61].

Here, we developed **S**ecure **Qu**ery Protocols for Genotype-Phenotype **D**atabases (**SQU**i**D**), a scalable framework designed to store and query genotype-phenotype databases in an ultra-secure cloud-based setting using homomorphic encryption. In our approach, we incorporate several key components: a ciphertext packing storage method to minimize the required storage space for encrypted data, a set of optimizations we developed to reduce query processing time, and an innovative cryptographic primitive (public key-switching) to enable homomorphic encryption for multiple users. We demonstrate that SQUiD is capable of efficiently executing various types of queries on large scale genotype-phenotype datasets, all the while maintaining the encryption of the data in the cloud. Specifically, it can perform tasks such as counting the number of patients in a filtered cohort, computing the Minor Allele Frequency (MAF) of genetic variants in a cohort, calculating Polygenic Risk Scores (PRS) for patients, and generating a cohort of genetically similar patients in remarkably short timeframes. Our findings highlight the potential of SQUiD as a valuable tool for secure, timely, and efficient analysis and interpretation of genetic and phenotypic data. At a time when data breaches are becoming increasingly common in healthcare settings, where data is a commodity, SQUiD provides a key resource to safeguarding patient privacy and enabling data providers to adhere to evolving laws and regulations, while ensuring the democratization of data.

## 3 Results

### Our conceptual framework allows ultra-secure interactions with encrypted genotype-phen-otype databases

Our conceptual framework is focused on solving real-world security challenges encountered in the storage and querying large-scale genotype-phenotype datasets. These challenges involve safeguarding the confidential information contained within such data from third party cloud providers and outside adversaries. Our framework is based on a scenario that involves three parties: the data owner, the researcher(s), and the public cloud. The data owner, who in many cases could be an organization such as the National Institutes of Health (NIH), possesses a vast amount of genotype-phenotype data that can be used for various analyses. Due to the large size of this data and the limited computing power and resources, the data owner encrypts the data and stores it in the public cloud. Their role is limited to authenticating clients who have permission to access the encrypted data, and they do not participate in the computation phase. The client, typically a researcher, seeks to perform computations on the data to obtain results. However, due to the large size of the data and the limitations of their computing power, it is not feasible for them to download, decrypt, and analyze the data locally. The client, therefore, interacts with the encrypted data deposited to the cloud. An overview of each parties role in the architecture of SQUiD is visualized in Figure 1. To ensure the privacy and security of sensitive data, the client must first obtain permission from the data owner to access the encrypted data. The cloud server does not have knowledge of the information contained in the data and, therefore, performs computations on the encrypted data using homomorphic encryption. This ensures that the sensitive information is protected while computations are being performed and the output is provided to the client in the encrypted form. Figure 2 describes the four key components of our framework: encrypted data storage, access authorization, query capabilities, and the API for user-friendly interactions with the database. Unfortunately, this scenario cannot be realized with traditional homomorphic encryption, which is based on a two-party (data owner and public cloud) system. In a traditional two-party system, the data owner encrypts the data with their public key and decrypts the results with their private key. Thus, the researcher cannot query and decrypt the results since they do not (and should not) have access to the data owner’s private key. To overcome this challenge, we adopted the concept of the Proxy Re-encryption system and developed its theoretical realization and practical implementation in homomorphic encryption, which we call a **public key-switching** technique [14,42,12,55].

**Figure 1:**
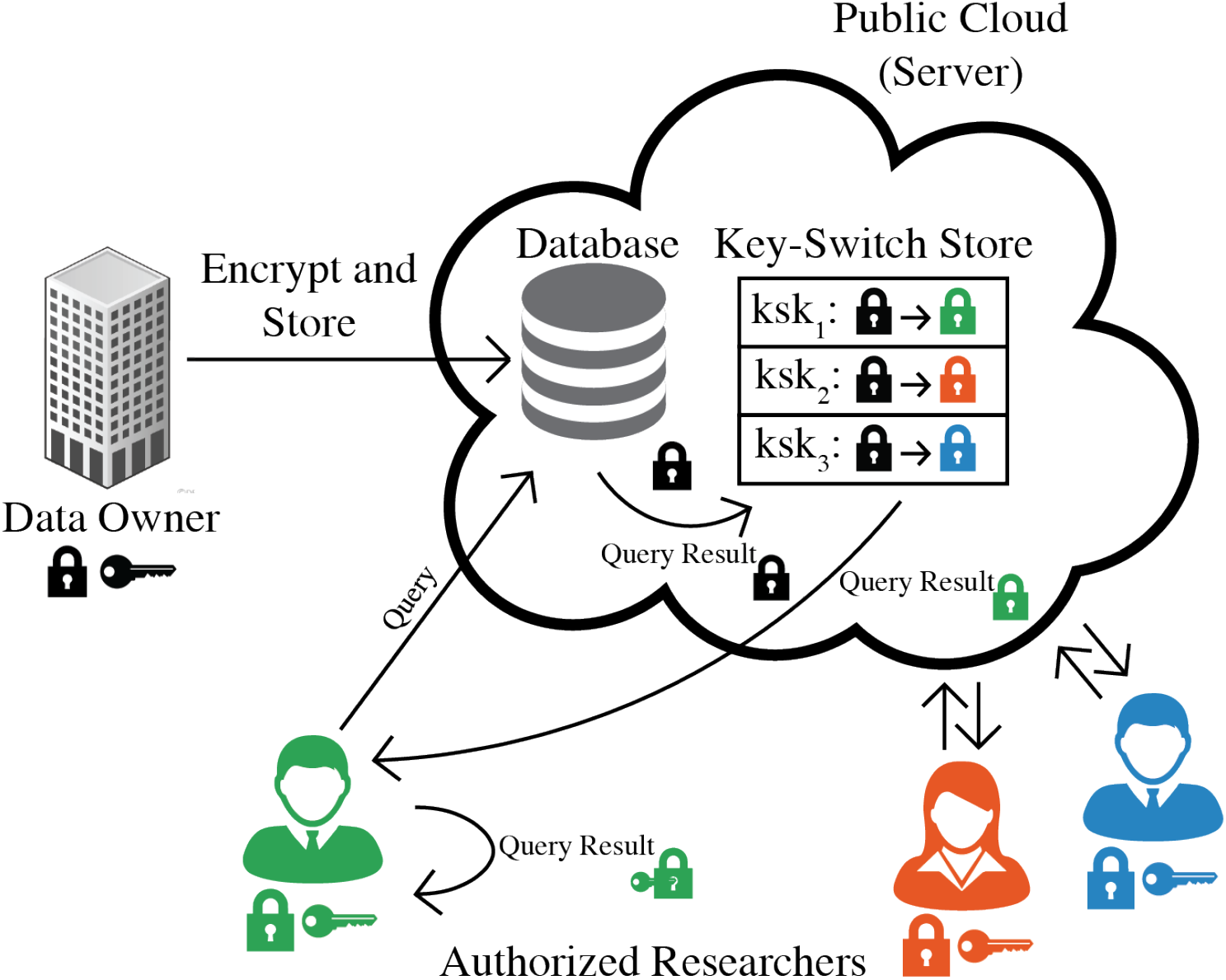
An architecture overview of SQUiD. Initially, the data owner uploads their encrypted genotype and phenotype to the public cloud. Within the cloud, only authorized researchers are permitted to securely query the data. Authorization is granted through possession of a key-switching key, which is stored in the key-switching store. When a researcher initiates a query on the databases, the database responds by encrypting the query result under the data owner’s public key. Subsequently, the key-switching store transforms this encrypted result to be under the querier’s key. The encrypted result is then sent back to the querier, who can decrypt it using their own secret key.

**Figure 2:**
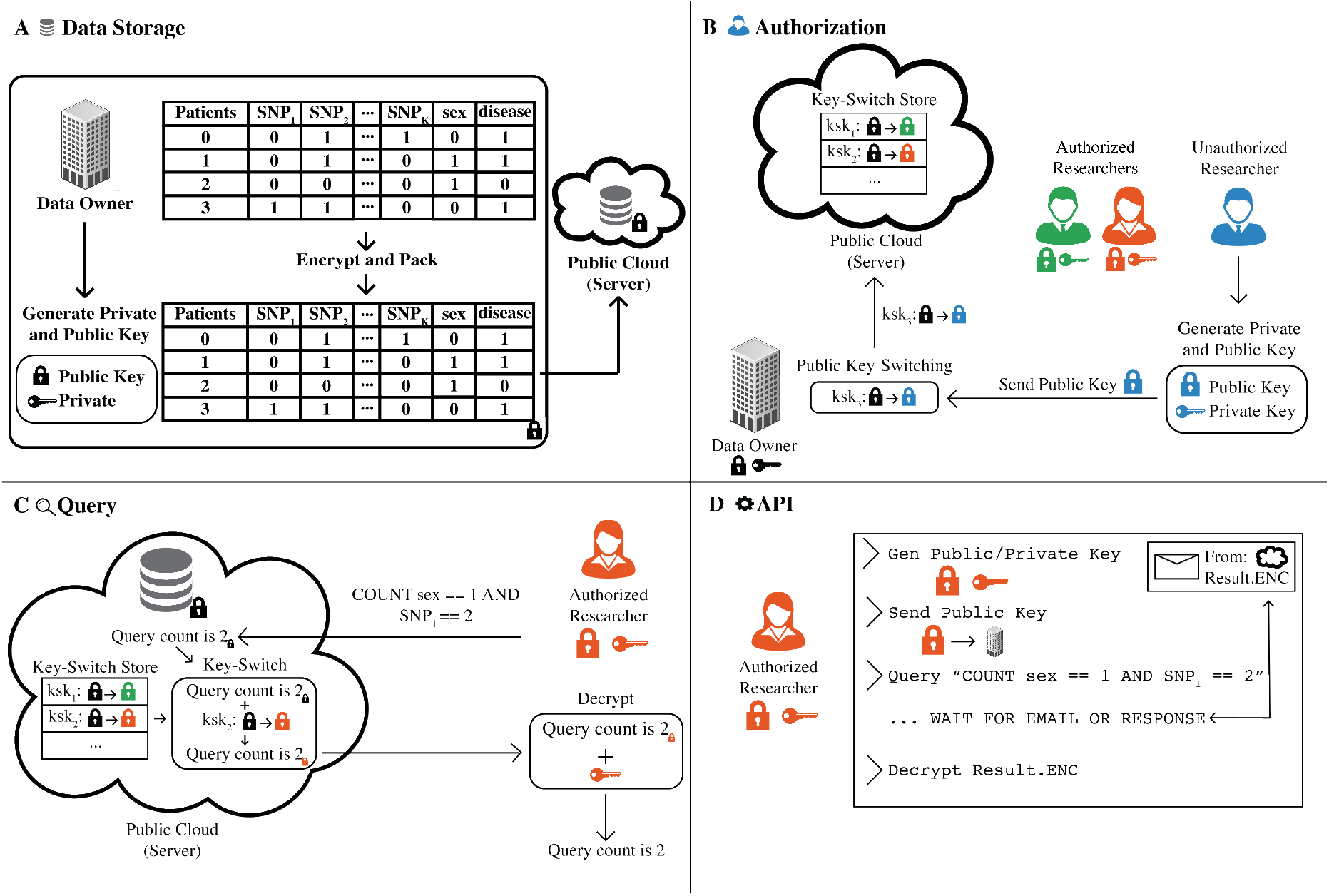
**(A) Data Storage**. The owner encrypts their data with a public key, then uses vertical packing to reduce storage costs before uploading the data to the public cloud. **(B) Authorization**. The onboarding process for a new researcher starts with the creation of their public and private key. The researcher sends their public key to the data owner for authorization. The owner authorizes the researcher by creating a key-switching key to switch the encryption of data to the researcher’s key, and uploading this key to the key-switching key store in the public cloud. The data owner can revoke a researcher’s access by removing the key-switching key from the store. **(C) Query**. An authorized researcher can submit one of four queries to the public cloud, which performs the necessary operations homomorphically on encrypted data under the data owner’s key. The result is then re-encrypted under the researcher’s public key and sent back for decryption. **(D) API**. We created a command-line API for researchers to use SQUiD easily. It generates a public and private key for the researcher, sends the public key to the data owner for authorization, sends queries to the server, and decrypts any encrypted results received via email or through a returned file.

The public key-switching operation serves as a cryptographic primitive facilitating the conversion of ciphertexts encrypted under one secret key to ciphertexts encrypted under a second secret key, without the need to decrypt the ciphertexts or possess access to the second secret key (see Methods section for the mathematical details). This capability holds immense value in establishing secure interactions with an encrypted database. For example, in SQUiD, the encrypted database can compute a researcher’s query under the encryption of the owner’s key, convert the computed result from an encryption under the owner’s key to under the public key of the researcher, and send this encrypted result to the researcher. Importantly, this conversion takes place without the need to decrypt the query result or disclose any information about it to the cloud. The researcher can effectuate this conversion by solely providing their public key, thereby circumventing any security risks associated with sharing their secret key.

When granting access to a new researcher, both the data owner and the researcher collaborate to create a public key-switching key, which is subsequently added to the key-switching store in the public cloud (Figure 2B). The key-switching store offers two significant advantages. Firstly, it allows the data owner to remain offline during any query, as the researcher exclusively interacts with the public cloud where the pre-calculated and stored public key-switching keys reside. Secondly, the data owner retains control over data access by managing the inclusion or exclusion of researchers’ public key-switching keys within the store, thus ensuring the ability to govern data access. We show that performance overhead from generating a key-switching key and key-switching a ciphertext under the encryption of one key to another to be less than a second (Supplementary Figure 1). The public key-switching key of each authorized researcher is stored in a dedicated key-switching store that dynamically expands to accommodate the number of authorized researchers. We show that the extra storage required for the key-switching store to be minimal (around 6.6 MB per researcher) (Supplementary Figure 2).

### Vertical packing allows efficient storage of encrypted large genotype-phenotype databases

We designed SQUiD to handle sensitive genotype-phenotype data from a large number of patients. SQUiD is specifically tailored to ingest data that has already undergone quality control and is stratified for population structure correction. Here we represent the data as a table with columns for basic attributes like age, sex, gender, etc; genotypes for single nucleotide polymorphisms (SNPs); and the phenotype or disease status of the patients, and rows for each patient in the database (Figure 2A). While each entry in the table needs to be an integer for the homomorphic encryption libraries we used, continuous phenotypes can be discretized into integers via scaling (see Supplementary Material for more details). The encryption of this data introduces additional storage requirements compared to its original unencrypted form. Consequently, the storage expenses associated with storing large genotype-phenotype databases in their encrypted state can be substantial. In order to optimize storage within the SQUiD framework, we adopt a vertical packing approach for our data organization (see Methods). This method involves storing the genotypes of multiple patients for a single SNP within a single ciphertext. By vertically packing our data (Figure 2A), we effectively reduce the number of ciphertexts necessary to accommodate a substantial volume of data, thereby minimizing the associated storage costs. Additionally, this approach enables us to perform high-throughput computations on the packed data efficiently, as operations involving packed ciphertexts, such as the addition of two packed elements, can be executed simultaneously (see Methods for details). Such packing still enables homomorphic updates (addition of new patients/SNPs/attributes) to the encrypted database without the need for decryption (see Methods for details).

We benchmarked the storage requirements of SQUiD on four different types of SNPs: ClinVar SNPs, Illumina Human1M-Duo v3.0 DNA Analysis BeadChip SNPs, Whole Exome Sequencing (WES) SNPs, and Whole Genome Sequencing (WGS) SNPs. These SNPs can be stored either at a per-chromosome level or genome-wide in SQUiD. Clinvar contains approximately 70,000 SNPs and Illumina BeadChip arrays contain approximately 1,072,820 SNPs. We estimated that around 8.2 million and 84 million SNPs would be observed in exomes and whole genomes at a population level, respectively, by using the data from 1000 Genomes Project [5]. We have benchmarked the packed storage of SQUiD against an unpacked homomorphic encryption storage, a storage encrypted with the industry standard AES-128-CBC, and a plaintext storage that stores SNPs as single bytes for the various SNP sets (Figure 3). We found that the storage cost for SQUiD is 16,384x better than the unpacked homomorphic storage cost (Figure 3), since a single ciphertext corresponds to data from up to 16,384 patients. Furthermore, vertical packing also allows the time for encryption to be reduced 16,384 fold compared to unpacked homomorphic encryption as fewer ciphertexts need to be encrypted (Supplementary Figure 3).

**Figure 3:**
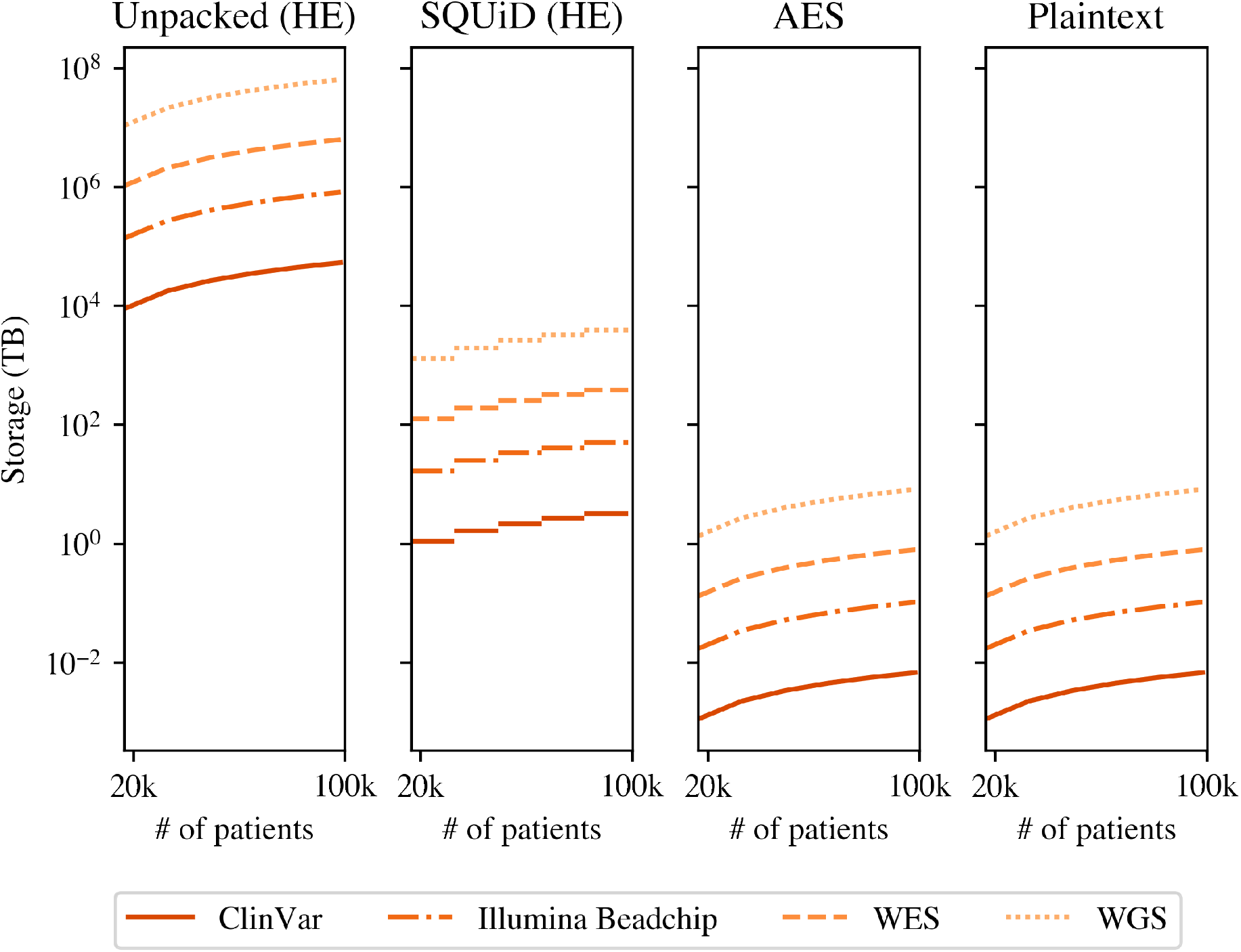
Plots showing the storage space required to store the ClinVar, Illumina Beadchip, WES, and WGS SNP genotypes with different schemes. The number of SNPs for WES and WGS is approximated using the 1000 Genomes Project.

### Enabling secure, scalable, and fast analysis of genotypes and phenotypes

We have devised four encrypted query functionalities within the SQUiD framework. This modular design allows for seam-less implementation of additional functionalities to accommodate diverse analysis requirements. Our queries include count, MAF, PRS and similarity. Figure 2C depicts how querying works under the public key-switching framework.

Count queries within the SQUiD framework ascertain the number of patients satisfying specific filters or equality checks. For example, a count query could count the number of patients with type-2 diabetes (T2D), whose SNP on gene TCF7L2 has a heterozygous alternative allele. MAF queries are employed to compute the MAF for a given target SNP within a filtered patient cohort. For instance, SQUiD can compute two MAF queries: one for a target SNP on the TCF7L2 gene within a cohort of patients with T2D, and another within a cohort of patients without T2D to study correlations between SNPs on the TCF7L2 gene and T2D. We can further add many different filters to build the cohort such as constraining it to patients with homozygous SNPs on a gene of interest. PRS queries involve the calculation and return of the PRS for all patients given a list of GWAS SNPs and their coefficients. PRS queries require only the coefficients and SNPs to be supplied post training such as those found on the PGS catalog [47]. Finally, similarity queries take a target patient’s encrypted genotype as input, build a cohort of genetically similar patients in the database through a scoring function like the squared euclidean distance, and output the number of similar patients with and without a particular disease of interest (see Methods and Supplementary Material). To evaluate the performance of each query, we conducted benchmarking against a plaintext implementation. The plaintext implementation keeps the genotype-phenotype data encrypted at rest (as mandated by the policies) and decrypts the necessary components of the data to compute the query in plaintext, while SQUiD keeps the data encrypted both at rest and during computation, enabling much stronger security as the data no longer has visibility to the computing party. This plaintext implementation models the current data access guidelines set by initiatives such as the dbGaP and UK Biobank where researchers download encrypted data, decrypt the data locally, and then analyze the data in plaintext [3].

On a dataset with 16,384 patients (*i.e*.,the maximum number of patients that can be stored in a single ciphertext) SQUiD can perform a count query with 16 filters in 15 seconds (compared to 0.013 seconds in plaintext), a MAF query with 16 filters in 20 seconds (compared to 0.027 seconds in plaintext), a PRS query with 1,024 SNPs in 5 seconds (compared to 0.8 seconds in plaintext), and a similarity query with 1,024 SNPs in 19 minutes (compared to 0.9 seconds in plaintext). We also show that our queries are easily parallelizable due to the linear nature of the queries. We benchmarked them in multi-thread environment and observed that the performance is significantly improved with more cores (Figure 5).

Our data show that all the functionalities implemented in SQUiD exhibit linear scaling relative to the size of their inputs. Specifically, the count and MAF queries scale linearly with the number of filters with a slope of 0.62, the PRS query scales linearly with the number of SNPs with a slope of 0.001, and the similarity query scales linearly with the number of SNPs given for the target patient with a slope of 0.068 (Figure 4). Our slopes consistently indicate a slow growth in runtime. Notably, the runtime of all protocols is proportional to the number of patients in the database and independent of the total number of SNPs in the database. A plaintext implementation of our protocols would also scale linearly with the number of patients in the database and the number of filters and SNPs involved in the query. Thus, SQUiD achieves optimal linear scaling as expected from a plaintext implementation, which signifies its ability to efficiently adapt to larger datasets in the future. Furthermore, with the expected decrease in the price of cloud computing in the future, the steady runtime observed for all queries ensures that increasing the size of the databases beyond the limits benchmarked in this study will yield steady performance outcomes, enabling real-world applications of SQUiD with biobank-scale data. We also show that the SQUiD’s communication cost for all queries except PRS query is constant regardless of the number of patients in the database while communication cost increases with the number of patients for all query types in plaintext (Supplementary Figure 4 and 5). Overall the communication is minimal. Comparable to an instagram post which has a maximum size of 4.3 MB (1,080 by 1,350 pixels) [1], most of our protocols use less than 50 MB.

**Figure 4:**
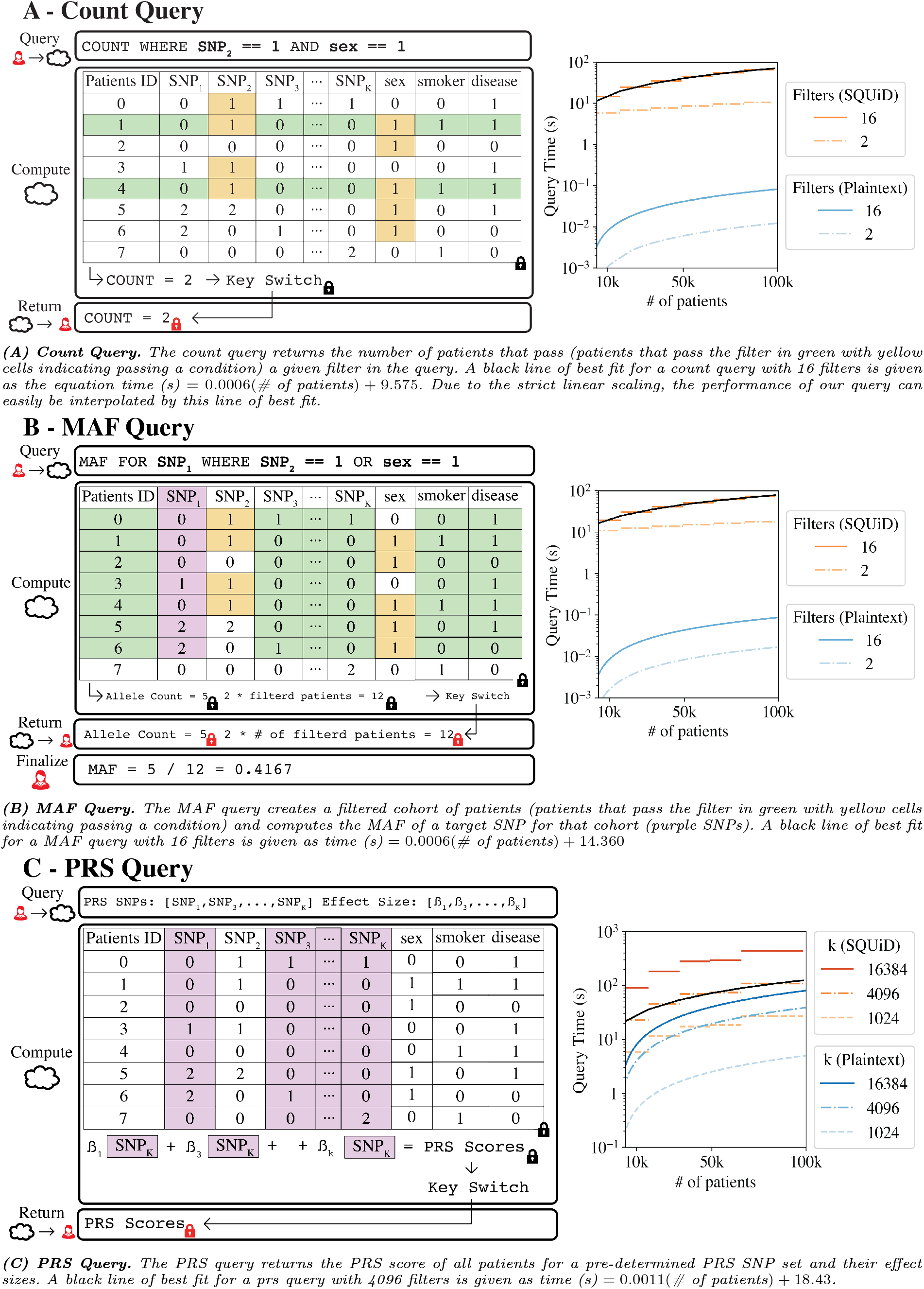
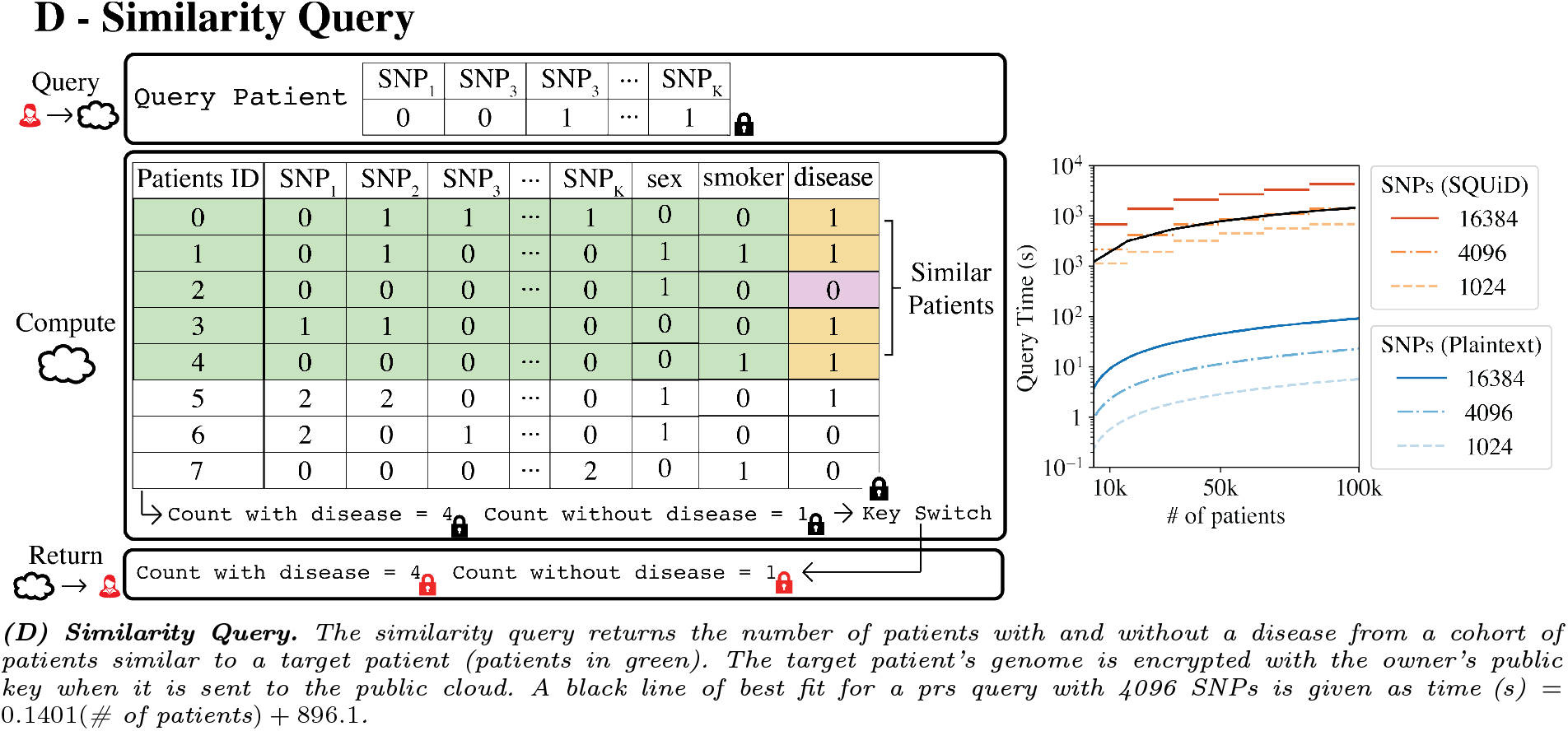
**(A, B, C, D)** For each query, the plots on the right show the end-to-end query time by varying the number of filters for the count and MAF query, by varying the number of SNPs and effect sizes (k) for the PRS query, and by varying the number of SNPs for the similarity query. The query time for SQUiD and the query time of a plaintext solution are shown for comparison. The plaintext solution works on a database encrypted with AES. For each plaintext query, the necessary components for the query are decrypted and then computed on.

**Figure 5:**
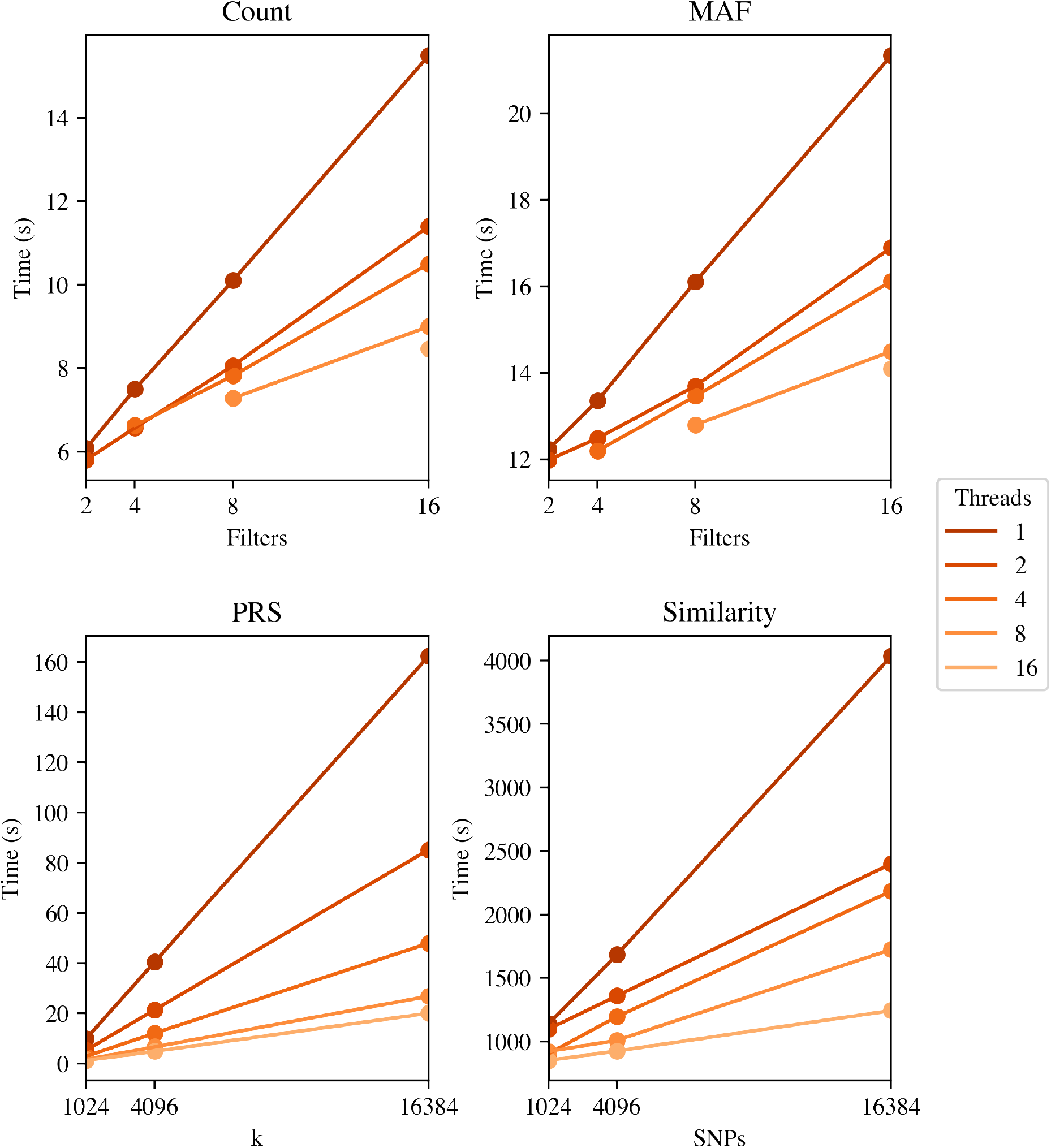
Plots of count, MAF, PRS, and similarity query time by the number filters, effect sizes (k), and SNPs in a varying the number of threads. We benchmarked the time for each query on a database with 16,384 patients using 2, 4, 8, and 16 filters for the count and MAF queries, and 1024, 4096, and 16384 SNPs for the PRS and similarity queries.

We also developed an API and a command line interface (CLI) to facilitate interaction with SQUiD, thereby enhancing its usability for researchers (Supplementary Figure 6). The API and CLI enable researchers to execute various queries and perform essential functions through simple commands. For instance, researchers can generate private and public keys required for encryption and authorization, send the public key to the data owner, execute all desired queries (See Supplementary Table 1 for API query parameters), and decrypt the returned query results. The data owner computes a public key-switching key, which is pushed to the cloud in the encrypted form. The API simplifies the deployment process for researchers who are not experts in privacy and security when utilizing SQUiD.

### SQUiD can reproduce known genotype-phenotype relationships in UK Biobank

We studied the relationship between patients with T2D and a control group in the UK Biobank dataset to assess the accuracy of the MAF and count queries in SQUiD. Firstly, we calculated the MAFs for the top five SNPs with the largest difference between T2D patients and the control group patients (Figure 6A). We compared the MAFs computed by SQUiD with the MAFs computed in plaintext to show there is no difference between them. Secondly, for these same five SNPs, we computed a chi-square statistic by using the allele counts for the control and case group (T2D in our case) [13]. We used the count query in SQUiD to get the allele counts and then computed the chi-square statistic in plaintext. The chi-square scores obtained from SQUiD queries are identical to the plaintext computation results (Figure 6B). Note that SQUiD does not directly execute GWAS, it has the capability to generate cohorts with specific attributes. We have shown that it can create accurate cohorts that will result in accurate GWAS demonstrated by the GWAS for T2D (Figure 6).

**Figure 6:**
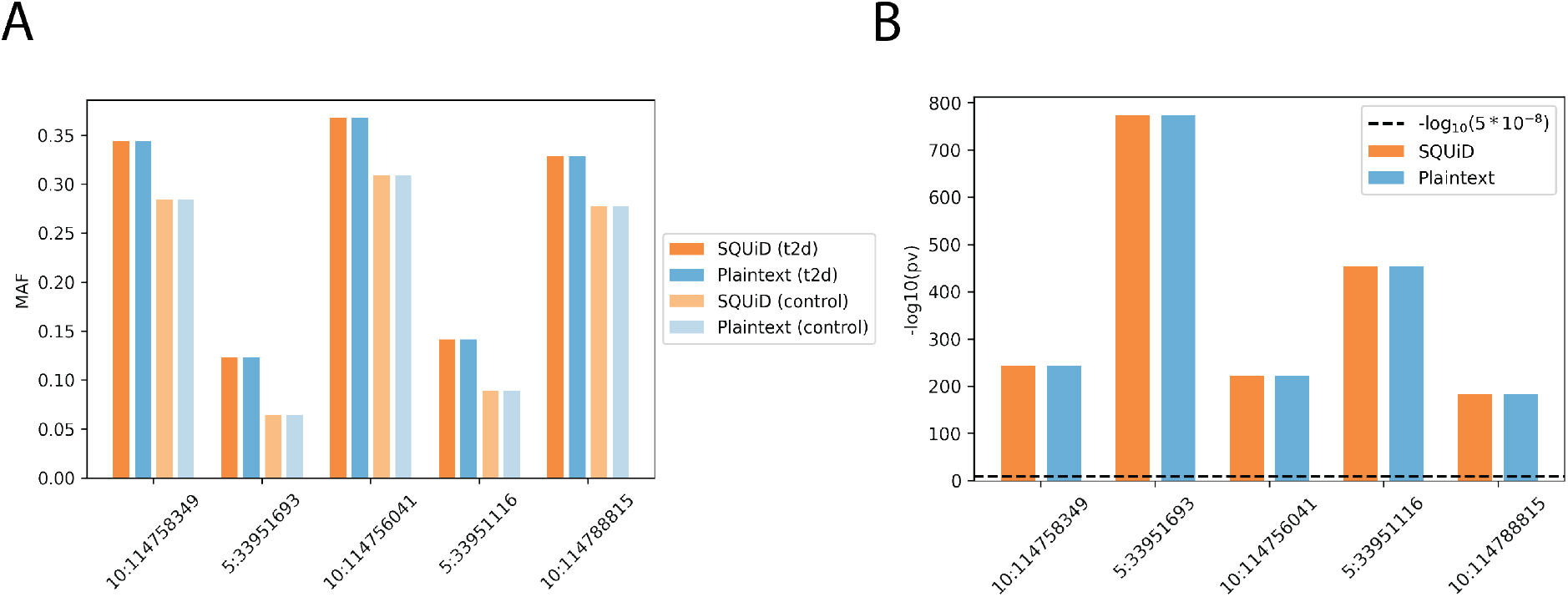
**(A)** Histograms of the MAFs of SNPs exhibiting the most substantial difference between control and T2D patient groups. The MAFs were calculated with SQUiD (orange), and in plaintext (blue). **(B)** Plot of -log(p-value) for the SNPs in (A).

We further evaluated the accuracy of SQUiD by replicating the sparse PRS calculations for standing height and T2D performed in the UK Biobank PRS study [65] using both plaintext calculations and the SQUiD PRS query. The standing height and T2D PRS use 51,209 and 183,830 SNPs, respectively.

They are the traits with the most number of SNPs involved in PRS calculations in the UK Biobank. We performed these calculations for 20,000 randomly selected patients in the UK Biobank. Our analysis revealed no observable difference in the PRS distribution and scores between plaintext and SQUiD queries (Figure 7). Notably, the sole discrepancy between the calculations arose from a marginal loss in precision. To accommodate the requirements of using integers in SNP effect sizes in SQUiD PRS queries, the effect sizes were multiplied by 100,000 and converted to integers. However, the resulting precision loss was minimal (Figure 7B,C).

**Figure 7:**
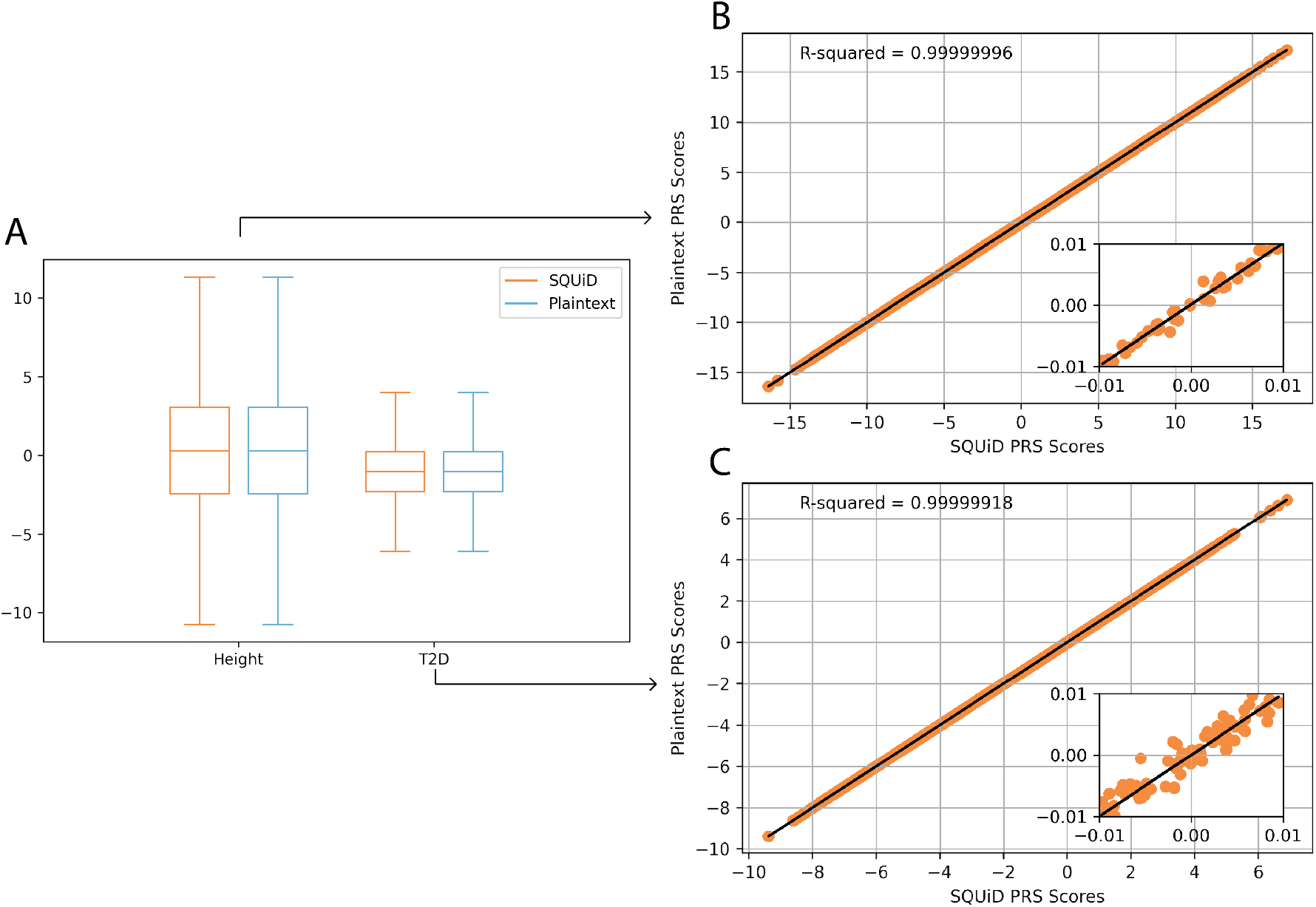
**(A)** Boxplots of the PRS score distributions of UK Biobank patients for standing height and type 2 diabetes (T2D) calculated with SQUiD (orange) vs plaintext (blue). **(B)** A scatter plot of the height PRS calculated by SQUiD vs. plaintext, where each point represents a patient. The black line is a line of best fit with an R^2^ of 0.99999996. **(C)** The same plot as (B) for T2D with an R^2^ value of 0.99999918.

## 4 Discussion

We introduce SQUiD, a novel, secure, and user-friendly queryable genotype-phenotype database implemented using homomorphic encryption. We envision SQUiD as a valuable tool for data owners, including hospitals, non-profit academic research institutions, and government health agencies, offering them a secure means to store genotype-phenotype data in the cloud while enabling authorized researchers to securely analyze this data. We propose that our system has the potential to replace existing genotype-phenotype databases, delivering enhanced security measures without compromising functionality. By employing homomorphic encryption, SQUiD offers a robust, scalable, and practical solution to mitigate privacy risks associated with sensitive genetic and phenotypic data. We demonstrate this by showing that SQUiD can scale with increasing numbers of patients and SNPs in a genotype-phenotype database, by performing a simple GWAS study on UKBB data, as well as by replicating PRS calculations in UKBB [65]. Note that we benchmarked SQUiD on a m5.8xlarge AWS instance with an Intel Xeon Platinum 8175 @ 3.1 GHz processor and 128 GB of memory. All our query protocols (count query, MAF query, PRS query, and similarity query) and encryption protocols (setup of the database) were run on single-threads unless otherwise indicated.

SQUiD leverages homomorphic encryption, which, to date, presented three key challenges. Firstly, traditional homomorphic encryption was designed for a two-party setting involving a server and a client. Secondly, it is known to incur a high storage cost. Lastly, analysis with homomorphic encryption tends to be slow. To overcome the two-party limitation, we developed a novel public key-switching operation. Note that this is significantly different than multi-key homomorphic encryption used in federated settings [52] as in multi-key homomorphic encryption, all parties own a subset of a single secret key. We anticipate that the public key-switching operation can be applied beyond our database design, offering broader applications in encrypted data sharing and encrypted computing that involve multiple independent parties. Furthermore, we demonstrate a significant improvement in storage efficiency through the application of a well-known vertical packing storage method, achieving a storage enhancement of 16,384 times compared to a naive homomorphic encryption solution. While storing SNP genotypes using homomorphic encryption increases storage costs relative to state-of-the-art encryption methods like AES, this approach is indispensable as homomorphic encryption allows execution of functions on encrypted data. Moreover, although our queries exhibit slower performance compared to plaintext solutions, we believe that the trade-off between security and performance is at acceptable levels. Moreover, multi-threading implementation improves the performance significantly. While plaintext queries can execute nearly instantly, our queries typically take seconds to minutes to complete. However, this performance overhead is unlikely to significantly impact the usability and utility of the framework for researchers. This is because the alternative is to download a large database and analyze the data locally, which is a much more time-consuming and resource-intensive process. Therefore, we believe that our framework offers a optimal balance of security and performance.

Although encrypted database systems do exist, to the best of our knowledge, none of them offer the same level of security guarantees and functionality as SQUiD. A developed secure database framework named CryptDB [56] offers efficient secure data storage and query performance. However, it does not offer the functionalities provided by SQUiD for two main reasons. Firstly, this framework is unable to compute the same set of queries as SQUiD. For instance, CryptDB lacks the ability to add *and* multiply encrypted database items, a necessary requirement for computing the linear combinations in PRS queries. Secondly, and more critically, CryptDB exhibits significant information leakage during equality checks used in the filtering process in count and MAF queries. Specifically, CryptDB exposes the count of unique items within the columns used for the equality checks. For genotype-phenotype databases that store SNPs with just three possible genotypes with known allele frequencies, CryptDB would expose the patients with the same genotypes for each SNP. This information could be combined with the known and well-studied population frequencies of each SNP to devise a simple attack that reconstructs the genotype values for each patient in the database, resulting in a complete breach of security. On the other hand, SQUiD offers a solution for issues of data protection, data privacy and stigma for researchers, funders, clinicians and patients. Furthermore the databases that keep data encrypted at rest with AES also do not provide the same security and functionality as SQUiD. For any query, these databases first need to decrypt the relevant data and then compute on them exposing the data to attacks. SQUiD can perform all queries without the need for decryption, significantly improving the security over existing systems.

Privacy-preserving MAF calculations using homomorphic encryption were proposed before [45]. Notably, SQUiD’s MAF query differs from this approach as it computes the MAF within a filtered patient cohort, where the filtering is done via protocols developed in this work. For a detailed mathematical exposition of these distinctions, refer to supplementary section 7.12.

We compared our patient similarity queries to existing private patient similarity queries (SPQ). Many existing SPQ protocols such as Wang et al. [68]privately compute patient similarity under the secure multiparty computation security assumptions, assuming non-colluding parties. Since SQUiD employs Homomorphic Encryption (HE), no assumptions about collusion between parties are necessary. Additionally, our query process involves a single round of communication, with the querying researcher sending a query to the cloud and receiving a prompt response. In contrast, the protocol outlined in [68] necessitates multiple rounds.

We also empirically compared SQUiD to [59] due to the similar security settings. The latter proposes a partial homomorphic encryption algorithm that supports only ciphertext addition and scalar multiplication operations for computing patient similarity using a squared L2-norm. We implemented the euclidean distance/squared L2-norm protocol [59] to the best of our understanding for comparison purposes. Supplementary Figure 7 shows that SQUiD can compute the squared L2-norm faster for larger datasets with an approximately 4x speed up for datasets with 50,000 patients.

We envision three use cases for this framework: 1-Funding agencies such as NIH can employ this framework to disseminate data currently available through the NIMH Data Archive (NDA) or Database of Genotypes and Phenotypes (dbGAP). 2-Multi-site consortia can employ this framework to disseminate data to their members while keeping the data secure in cloud storage. 3-Learning health systems can employ this framework to disseminate data to their researchers while keeping the data secure in cloud storage. Our secure framework is designed to enable users to form specific patient cohorts based on desired characteristics. Within this system, users can also determine the distribution of PRS for a particular disease across various patient populations. For example, one can explore the PRS distribution for schizophrenia among patients diagnosed with bipolar disorder. Additionally, the framework allows for the analysis of disease outcomes in patients who share genetic similarities with a specific patient of interest, facilitating more personalized and targeted approaches to healthcare and research.

In conclusion, SQUiD presents an innovative and impactful solution for a world grappling with escalating concerns surrounding security and privacy of genetic and clinical data. By circumventing the challenges posed by the ever-changing, heterogeneous landscape of data protection laws, SQUiD offers a robust framework to safeguard sensitive information. Moreover, we firmly believe that SQUiD has the potential to enhance patient trust by ensuring the security and controlled utilization of their data for specific research purposes, thus has the potential to increase participation in genetic research. Lastly, although this study focused on genotype-phenotype analyses for proof of principle, SQUiD’s modular design allows for the integration of other data modalities and analytic approaches, as the need arises.

This adaptability will be critical at a time when precision medicine research is rapidly expanding to encompass more complex molecular and clinical datasets.

## 5 Code Availability

The code for SQUiD, the SQUiD API, and the SQUiD CLI is available on GitHub for non-commercial use at https://github.com/G2Lab/SQUiD/.

## 6 Acknowledgments

This study is funded by NIH grants R00HG010909 and R35GM147004 to GG, a Warren Alpert Foundation grant to GG and TL, and a gift from Woodnext Foundation to TL.

## 7 Methods

### 7.1 Security and threat models

Our security assumption is based on the current data-sharing policies within many public and private entities. That is, the data owner and authorized researchers are mutually trusted. Thus, authorized researchers are allowed to query the genotype-phenotype data that do not threaten the confidentiality of patients according to the data use agreements. The inherent data leakage from query results and potential inference attacks from authorized researchers are therefore not considered.

Meanwhile, genotypes and phenotypes as well as a subset of the queries are protected from the public cloud and attackers. More precisely, we consider the following three threat models for database management [35,10]:

− Snapshot attackers that obtain a snapshot of the database
− Persistent passive attackers that compromise the cloud server to obtain not only the database but also queries and all server’s operations
− Active attackers that fully compromise the server to deviate from pre-designed protocols for queries In our SQUiD construction, snapshot attackers receive ciphertexts of the BGV homomorphic encryption scheme. We follow the Homomorphic Encryption Standard [8] to choose BGV parameters that provide a 128-bit security level against known attacks. Consequently, the security towards snapshot attackers inherits from BGV’s IND-CPA security, *i.e*., the ciphertexts are almost indistinguishable from random characters.

For persistent passive attackers, there are many ways that querying encrypted databases can result in private information leakage [29,41,49,53]. Most prominent ones include leakage through (1) *access pattern*, which determines if certain records are consistently accessed; and (2) *search pattern*, which indicates if and when an encrypted query is repeated. Many cryptosystems, including property-preserving encryption (PPE) [7,48] and searchable encryption (SE) [26,64], fail to protect against these types of information leaks. This is primarily due to their inherent functionality, which inadvertently discloses properties of datasets, thereby compromising privacy. However, homomorphic encryption (HE) schemes such as BGV provide a solution that does not leak access and search patterns [43]. Using HE to encrypt databases propels algorithms that have to touch all the relevant records in the dataset for a single query. For example, to find out whether an encrypted input is in the encrypted database, the input needs to be compared with every single encrypted value in the database homomorphically. This prevents access pattern leakages since the access pattern remains uniform for all queries. In addition, search pattern leakages are prevented due to the IND-CPA security under carefully selected parameters, since encrypted queries are indistinguishable from one another, regardless of their contents [43].

It is worth mentioning that persistent passive attackers do not learn additional information about the database from knowledge of the server’s computation patterns. Precisely, when an authorized researcher sends a query *f*, the server performs a series of operations on the encrypted database **Enc**(*m*) to obtain **Enc**(*f* (*m*)). The function *f* is in plaintext for Count, MAF and PRS queries and contains ciphertexts for similarity queries. In all these cases, the computation pattern for the server is predefined and contains operations such as homomorphic additions, multiplications, and key switching. As such, inference attacks from persistent passive attackers are also prevented, as only computational patterns of different functions are revealed but not any computation result *f* (*m*).

While the problem of defending against active attackers is challenging and still unsolved [34,35], our SQUiD construction provides reasonable mitigation towards active attackers. Namely, active attackers can deviate from pre-determined operations in SQUiD and therefore send wrong computation results to authorized researchers, but they can not learn information about the database.

### 7.2 Homomorphic Encryption

Encryption is a procedure that maps the *plaintext* data into its *ciphertext*, such that the plaintext can not be deduced from the ciphertext without knowing the secret key. Homomorphic encryption (HE) is a class of encryption schemes with an additional property: computations can be performed over ciphertexts without knowing the secret key. Figure 8 visualizes this property in a commutative diagram.

**Figure 8:**
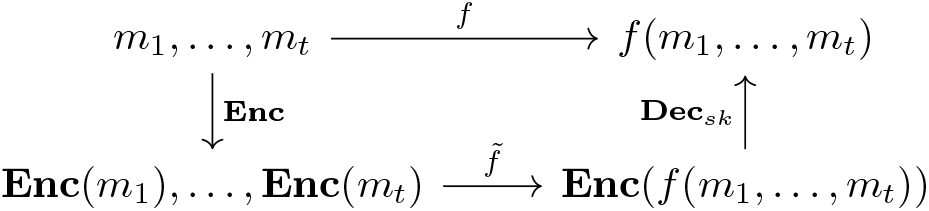
The homomorphic evaluation of a function f on ciphertexts

HE ciphertexts contain a *noise* component, whose value grows with homomorphic operations. This is controlled by pre-fixed HE parameters, which is also used to set a noise budget. If the number of operations in an algorithm is too large such that the noise consumption exceeds the budget, then the result can no longer be decrypted correctly. To avoid this, a *bootstrapping* operation is introduced to refresh the ciphertexts, enabling the *fully* homomorphic encryption (FHE) schemes that support evaluations of *arbitrary* circuits for different operations including multiplications and additions (*i.e*., arbitrary *f*) [31]. Detailed realizations of homomorphic operations are included in the supplementary material.

### 7.3 Brakerski-Gentry-Vaikuntanathan scheme

The Brakerski-Gentry-Vaikuntanathan (BGV) scheme is an FHE scheme that relies on the hardness of the Ring Learning-with-error (RLWE) problem [51]. Its basic building blocks are homomorphic addition ADD and multiplication MULT. Since any computable function can be realized with additions and multiplications, the homomorphic evaluation of any computable *f* can be realized with ADDs and MULTs. Bootstrapping in BGV is a very costly operation [22,39]. It is, therefore, common to use BGV in the *levelled* manner, *i.e*., to choose the HE noise parameter with large noise capacity such that computations can be performed without bootstrapping. Our study uses the levelled version of BGV.

BGV allows efficient computations in the amortized sense. It supports Single Instruction Multiple Data (SIMD) operations, which allows multiple values to be packed into one BGV ciphertext, enabling computations over a single ciphertext to be performed on all packed values in an efficient manner [63]. Details of the SIMD packing are included in the supplementary material.

### 7.4 Public key-switching

In general, HE binary operations only support input ciphertexts that are encrypted under the same key. Therefore, in the scenario of multiple users each holding their own keys, there is a natural need to convert a ciphertext encrypted under one key to another ciphertext that encrypts the same message under a different key. A naive approach is to decrypt and re-encrypt with a different key, but this exposes the original secret key and the message to the party that performs this procedure. To prevent such leakages, the above procedure can be done *homomorphically* such that the evaluation party can not access the message in the clear. Such a technique is called *key switching*. Mathematically, when converting the key system from (`pk, sk`) to (`pk`^*a*^*st*, `sk`^*^), the evaluation party does not need to know `sk`, but a key-switching key ksk_(sk→sk*)_ which leaks no information about secret keys.

Our scenario exploits the key-switching key ksk_(sk→sk*)_. While the traditional key-switching key generation uses both `sk` and `sk*`, only `sk` and `pk*` are needed in our design, hence it is called public key-switching. This design preserves the confidentiality of `sk*` as it does not need to be shared to compute the key-switching key. In our scenario, the secret key of the authorized researchers does not need to be sent to the data owner to generate the key-switching key. Please see supplementary material for the mathematical details of the realization of public key-switching with BGV and how we control the increasing noise.

### 7.5 Database construction with vertical packing

The dataset in SQUiD is represented as a matrix *M* = {*m*_(*i,j*)_|1 *≤ i ≤ r*, 1 *≤ j ≤ k*}, where *r* is the number of patients, *k* is the number of attributes (features), and the value in position (*i, j*) corresponds to the *j*-th feature of the *i*-th patient (*e.g*., the genotype of *j*-th SNP of *i*-th patient). We use the term *vertical* or *horizonal* for the direction in the matrix, which corresponds to a feature for all patients or attributes, respectively.

As we explained earlier, BGV supports packing multiple messages into one ciphertext. SQUiD packs elements *vertically* : let *𝓁* denote the packing capacity in a ciphertext, then the *r* elements in the *j*th column are encrypted into ⌈*r/𝓁*⌉ ciphertexts

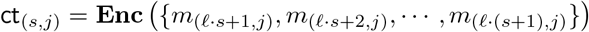

where 1 *≤ s ≤* ⌈*r/𝓁*⌉ and *m*_*i,j*_ is considered as 0 for *i > r*. Overall, entire dataset is encrypted into *C* = {`ct`_(*s,j*)_ | 1 *≤ s ≤* ⌈*r/𝓁*⌉, 1 *≤ j ≤ k*}.

The update, insert, and delete operations on a vertically packed encrypted database vary slightly from their typical implementations.

− *Update:* To update a single value *m*^*′*^ at index *i, j*, a new encryption of `ct`_(*s,j*)_ where *s* = ⌈*i/𝓁*⌉ needs to be uploaded where

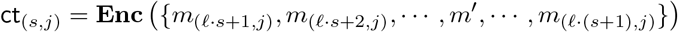
− *Insert:* To insert a new row at *r* + 1, if ciphertexts are not fully packed (i.e., *𝓁* ∤ *r*), then the last row of packed ciphertexts contains zeros at row index *r* + 1, which are updated. Otherwise, the following *k* fresh ciphertexts are added, forming the last row of *C*.

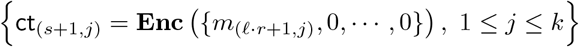
− *Delete:* To delete an entry at index *i, j*, a plaintext, which encodes zero at the *i* mod *𝓁*-th slot and one elsewhere is multiplied with ct_(⌈*i/𝓁*⌉,*j*)_.

Note that update and insert operations both upload new ciphertexts with low noise, but the delete operation increases the noise with a plaintext-ciphertext multiplication. To bound the noise growth, we set a number *α* for the maximum times of consecutive delete operations. On the (*α* + 1)-th time to delete an entry, an update should be performed instead, after which *α* deletes are again allowed. For SQUID with our experimental parameters, the value *α* is taken to be 1.

### 7.6 Functionalities

In this section, we describe the supported functionalities of SQUiD and the evaluation procedures using homomorphic encryption.

#### Count queries

The first category of queries is to *count* the number of patients *whose* attributes satisfy certain conjunctive (`AND`) and/or disjunctive (`OR`) relations. Its evaluation contains two stages, filtering and vertical aggregation.

*Filtering* Suppose the researcher specifies *τ >* 1 selection criteria (either in plaintext or ciphertext) and their relation (`AND` and/or `OR`). The filtering stage outputs a *predicate vector* **p** composed of *r* encrypted binary numbers. If the element **p**[*i*] decrypts to 1, then the patient *i* is in this pre-defined cohort.

First, we explain how to homomorphically check a single selection criterion, which amounts to performing a homomorphic equality test between the given value in a query and a value in the matrix. The key idea is to find a polynomial representation, which can be evaluated as a sequence of homomorphic additions and multiplications.

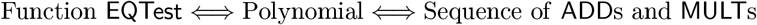

Without loss of generality, we consider the inputs of `EQTest` as genotype values in {0, 1, 2}, and denote them as *u* and *v*. As shown in Table 1, this function determines a unique truth table.

**Table 1:** The truth table of EQtest(u, v) for SNPs.

| $\begin{array}{c\vert c} & u \\ \hline v & \end{array}$ | 0 | 1 | 2 |
| --- | --- | --- | --- |
| 0 | 1 | 0 | 0 |
| 1 | 0 | 1 | 0 |
| 2 | 0 | 0 | 1 |

We derive the polynomial representation of `EQTest`(*u, v*) as follows. Let *v* be an encrypted matrix value, and *u* be the query value, which can be either in the clear or encrypted depending on the researcher. If *u* is provided in the clear, then we can interpolate the *u*-th column of the truth table 1 into a degree-2 polynomial *F*_*u*_ with input variable *v*. If *u* is also encrypted, then we precompute a polynomial *F* of degree 4 that maps 0 to 1 and {*±*1, *±*2} to 0, whose input variable is *u* − *v* ∈ {0, *±*1, *±*2}.

Second, we explain how to homomorphically combine the results of multiple equality checks using `AND` and `OR`. Let 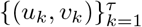 be the set of (encrypted or unencrypted) queries and (encrypted) matrix values, then for each patient *i* we compute the expression homomorphically as Equation (1), where *d* and *b* are constants in Table 2. The evaluation decrypts to 1 if the data of the patient *i* matches the selecting criteria, and 0 otherwise.

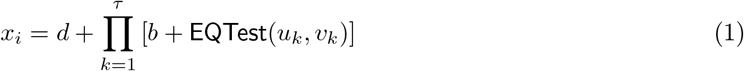

**Table 2:**
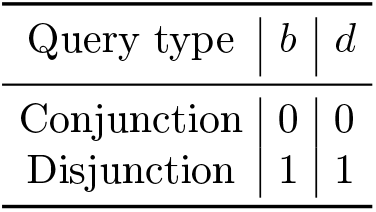
Constants in circuit (1) [46].

| Query type | $b$ | $d$ |
| --- | --- | --- |
| Conjunction | 0 | 0 |
| Disjunction | 1 | 1 |

*Vertical Aggregation* Suppose each ciphertext provides *𝓁* SIMD slots, then the predicate vector for *r* patients is batched into ⌈*r/𝓁*⌉ ciphertexts. The procedure of summing over these batched messages is a *vertical* aggregation.

Our design fully exploits the advantages of parallel computing. Namely, we perform *𝒪* (*r/𝓁*) homomorphic additions with additive depth *𝒪* (log (*r/𝓁*)) to obtain one ciphertext, whose *𝓁* components are then aggregated with *𝒪* (log *𝓁*) homomorphic rotations and additions. Please see supplementary material for details of homomorphic addition and rotations with BGV.

#### PRS queries

The second category of queries is to obtain *polygenic risk scores* of all the patients.

##### Definition 1.

*The polygenic risk score (PRS) of a patient is a linear combination of values of attributes in a subset S. For given coefficients (i.e*.,*effect sizes)* {*β*_*j*_}_*j*∈*S*_, *the PRS for patient i is* 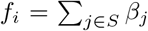, *where m*_(*i,j*)_ *is the genotype of the j-th SNP for i-th patient*.

The PRS for each patient can be calculated with homomorphic multiplication and additions. Please see supplementary material for details of homomorphic addition and multiplications with BGV.

*Horizontal aggregation* PRS queries aggregate information horizontally. We use parallel computing to minimize the execution time, and as can be seen from the Results section, answering PRS queries is relatively fast.

#### MAF queries

The third category of queries is to calculate the *minor allele frequency* for a target SNP of a filtered cohort of patients.

##### Definition 2.

*Minor allele frequency (MAF) is the frequency at which the minor allele occurs in a given population or a cohort. Let* **p** *be the predicate vector for r patients, where* **p**[*i*] *indicates whether the patient i is in the cohort. Then, for the dataset M* = {*m*_(*i,j*)_}, *the MAF for SNP j with* **p** *is*

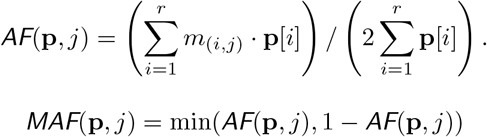

As the homomorphic division and minimum comparisons are currently expensive operations, the cloud instead computes the numerator and denominator homomorphically and then returns the results to the clients for decryption, division, and the minumum operation.

The plaintext modulus can be adjusted to be as low as the number of patients in the database, the MAF query may produce the numerator distributed across multiple slots if twice the number of patients exceed the plaintext modulus. These slots adding up to the overall numerator value, and it is the responsibility of the client to aggregate these values, calculating the final numerator. The subsequent steps for finalizing M AF c alculations remain unchanged.

#### Similarity queries

The fourth category of queries determines whether a specific individual (denoted as *d*) is genetically similar to patients *with* a certain disease or those *without*. There are two similarity metrics for researchers to choose from.

##### Definition 3.

*Suppose the database storesk attributes and the last attribute is the disease*.

1. *The L*^2^*-distance similarity score* 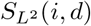 *is defined as*

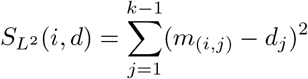
2. *The Jaccard similarity score S*_*Jcd*_(*i, d*) *is defined as*

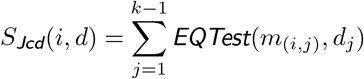

*where `EQTest`*(·, ·) *equals to 1 if two inputs are equal and 0 otherwise*.

In other words, the similarity score *S*_(·)_(*i, d*) horizontally aggregates the result of the squared difference or `EQTest`.

As a result of this query, the researcher will receive three encrypted values *r*_1_, *r*_2_, *r*_3_ from the cloud. The value *r*_1_ is the number of patients with this disease, *r*_2_ is the number of patients that are genetically similar to the target *d*, and *r*_3_ is the number of patients with this disease that are similar to the target *d*. After decryption, the researcher compares the ratio *r*_3_*/r*_1_ with (*r*_2_ − *r*_3_)*/*(*r* − *r*_1_) and determines whether the individuals similar to target individual *d* are more likely to have the disease.

These three values are homomorphically computed as follows.

1. Similar to the filtering method in Section 7.6, the cloud computes a predicate **p** for patients with this disease, whose vertical aggregation gives *r*_1_.
2. To count similar patients, the cloud computes the similarity score *S*_(*·*)_(*i, d*) between the target *d* and patient *i*. Then the cloud homomorphically checks if *S*_(*·*)_(*i, d*) is greater than the pre-determined threshold *t*, which is done by evaluating the interpolation polynomial of degree Range(*S*_(*·*)_(*i, d*))−1. In our implementation, We use the Paterson-Stockmeyer method [54] to evaluate polynomials efficiently. As such, we get a predicate **p**_*s*_ whose vertical aggregation gives *r*_2_.
3. Multiplying the two predicates **p** and **p**_*s*_ component-wise realizes the `AND` relation and leads to another vector, whose vertical aggregation gives *r*_3_.

## Supplementary Information

### 7.7 Security of Homomorphic Encryption and RLWE

Most widely-used homomorphic encryption schemes [17,16,28,23], including the BGV scheme we employed, rely on the hardness of a mathematical problem called the Ring Learning With Errors (RLWE) problem, *i.e*., the ring version of Learning with Errors (LWE) problem. The hardness of the LWE problem has served as a common assumption due to its quantum reduction to a hard lattice problem proposed by Regev [57] and its robustness against various attacks [15,11]. Later, Lyubashevsky *et al*. [51] introduced the RLWE problem and demonstrated its reduction to the worst-case lattice problems over ideal lattices.

Compared to LWE-based cryptographic schemes, RLWE-based schemes are more efficient, especially for power-of-2 cyclotomic rings [58].

To delve deeper into the RLWE problem, we introduce the following notation: *Φ*_2*n*_(*X*) represents the 2*n*-th cyclotomic polynomial, an integer-coefficient polynomial ring *ℛ* = Z[*X*]*/Φ*_*m*_(*X*) where *Φ*_*m*_(*X*) represents the *m*-th cyclotomic polynomial. a modulus *Q*, an error distribution *χ*, and ← for a uniform sampling.

Given a secret *s* ∈ *ℛ*_*Q*_ where *ℛ*_*Q*_ = *ℛ/Qℛ*, the hardness of RLWE guarantees that the following two distributions in *ℛ*_*Q*_ *× ℛ*_*Q*_ are (computationally) indistinguishable:

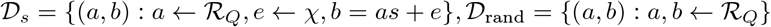

As such, the secret *s* is masked using a pair of elements, which are indistinguishable from two uniform elements. Parameter choices of RLWE-based HE schemes depend on the state-of-the-art attacks on the LWE problem. Concrete parameters are generally estimated following [9], which are also standardized in the white paper [8].

### 7.8 Experimental Setup and HE parameters

We benchmarked SQUiD on a c4.8xlarge AWS instance with an Intel Xeon E5-2666 v3 @ 2.9 GHz processor and 60 GB of memory. All our query protocols (count query, MAF query, PRS query, and similarity query) and encryption protocols (setup of the database) were run on single-threads.

For all our benchmarking, we employed parameters conforming to the 128-bit security standards set by the Homomorphic Encryption Standard [8]. Specifically, we set *n* to 32,768, *logq* to 880, and *p* to 131,071, where *n* is the dimension, *q* is the ciphertext modulus, and *p* is the plaintext modulus. With these parameters, the slot size is 16,384, enabling us to store up to 16,384 patients in a single ciphertext.

### 7.9 API Parameters for Queries

#### Count

Our simplest query counts the number of patients who pass a filter. This filter can be either conjunctive (And) or disjunctive (Or) and consist of only equality causes. For example in figure 3, we have the query “COUNT WHERE SNP1 == 1 and sex = 1” which will count the number of female patients where the first SNP is 1. We compute this query by first computing a predicate for each patient if they pass the filter. Since homomorphic encryption only supports addition and multiplication, we compute the filter consisting of AND gates, OR gates, and equality function using polynomial approximations. The domain of the inputs to all these functions is limited to (0,1,2) for SNPs and (0,1) for all other variables in our database. Thus, the degree of these polynomials is small which keeps SQUiD performant.

#### MAF

Our minor allele frequency (MAF) query computes the MAF of a target SNP on a filtered cohort of patients filtered using a similar filtering technique from the count query. Our MAF queries return two values, the allele count and the number of filtered patients times two. The researcher has to finalize the MAF computation by dividing the returned allele count by the returned number of filtered patients times two. We move the division operation to the research because division is an expensive operation to perform homomorphically.

**Supplementary Table 1:** List of API parameters for count, MAF, PRS, and similarity queries with examples.

| Query | Parameters | Query |
| --- | --- | --- |
| Count | Filter: array of pairs of ints<br>Conjunctive: boolean | Filter is $[(1,1),(3,1),(4,0)]$<br>Conjunctive is 1<br>Returns the number of patients in the database that have a first SNP with value 1, third SNP with value 1, and fourth SNP with value 0. |
| MAF | Filter: array of pairs of ints<br>Conjunctive: boolean<br>Target SNP: int | Filter is $[(1,1),(3,1),(4,0)]$<br>Conjunctive is 0<br>Target SNP is 2<br>Returns the MAF for SNP 2 of patients in the database that have a first SNP with value 1, third SNP with value 1, or fourth SNP with value 0. |
| PRS | PRS: array of pairs of ints | Parameters are $[(1, 4), (2, 10), (5, -4)]$<br>Returns the PRS score for each patient by summing up the multiplication of the first SNP by 4, the second SNP by 10, and the fifth SNP by -4 |
| Similarity | Target patient: ciphertext<br>Threshold: int<br>Disease column index: int | Target patient is MO8v3J7kJt (Encrypted)<br>Threshold is 10<br>Disease column index is 100<br>Returns the number of patients with and without a disease at column index 100 who are similar to the target patient by a threshold of 10 using a L2 similarity score. |

#### PRS

Our polygenic risk score (PRS) query computes a PRS given a set of SNPs and a set of effect sizes. A PRS score is calculated with a linear combination of the SNP and the effect sizes which we can naturally compute using homomorphic encryption. The effect sizes, which are often represented as floating point numbers, are scaled to integers within SQUiD as SQUiD only supports integer inputs. The resulting scores are scaled down accordingly.

#### Similarity

The similarity query counts the number of patients with and without a disease from a cohort consisting of patients similar to the target query patient. This target patient is encrypted with the data owner’s public key before it is sent to the cloud to protect the target patient data. To compute the similarity query, we first compute a similarity cohort by scoring the similarity of the target patient with every patient in the database. We score patients based on their squared Euclidean distance (squared L2 distance) from the target patient. If these scores are beneath a predetermined threshold, the patient is considered similar to the target patient and added to the similarity cohort. Next, we count the number of patients with and without a disease in this cohort.

### 7.10 The BGV scheme

Brakerski-Gentry-Vaikuntanathan (BGV) [17] is a well-known lattice-based homomorphic encryption scheme that allows for computations over encrypted data. Its lattice-based structure further provides reasonable quantum resistance. Realizing computations as sequences of additions and multiplications, BGV provides a large computation capacity with reasonable parameter choices.

Furthermore, it supports computations in a SIMD manner, which significantly enhances the amortized performance.

Below we describe the basic procedures in BGV, the realization of public key-switching and its noise control. Mathematical and cryptographic notations are listed in Table 2.

**Supplementary Table 2:** Notations of BGV homomorphic encryption scheme.

|  |  |
| --- | --- |
| <u>Mathematical Basics:</u> |  |
| $t$ | plaintext modulus |
| $Q$ | ciphertext modulus |
| $\mathbb{Z}$ | set of integers |
| $\mathbb{Z}_t$ | the ring of integers modulo $t$, i.e., $\mathbb{Z}_t = \mathbb{Z}/t\mathbb{Z}$ |
| $\mathbb{Z}_Q$ | the ring of integers modulo $Q$, i.e., $\mathbb{Z}_Q = \mathbb{Z}/Q\mathbb{Z}$ |
| $X$ | variable of polynomial |
| $R$ | the cyclotomic ring $\mathbb{Z}[X]/(\Phi_m(X))$ where $\Phi_m(X)$ is the $m$-th cyclotomic polynomial. The degree of $\Phi_m(X)$ is $n = \phi(m)$, where $\phi(\cdot)$ denotes the Euler totient function. |
| $R_t$ | $\mathbb{Z}_t[X]/\Phi_m(X)$ |
| $R_Q$ | $\mathbb{Z}_Q[X]/\Phi_m(X)$ |
| <u>HE Parameters:</u> |  |
| $\chi_{\text{ter}}$ | uniform ternary distribution with coefficients from $\{-1, 0, 1\}$ |
| $\chi_{\text{err}}$ | error distribution of scheme, typically a discrete Gaussian distribution unless specified otherwise |

#### Key Generation, Encryption and Decryption

Sample the secret key `sk` from *χ*_ter_, and the public key is computed as follows

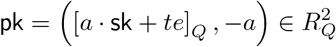

where *a* ← *u*_*Q*_ and *e* ← *χ*_err_. Anyone can see the public key, but this leaks no information on the secret key `sk`, as guaranteed by the hardness of the ring learning-with-errors (RLWE) problem.

With the public key, anyone can generate a ciphertext for a plaintext *m* by computing

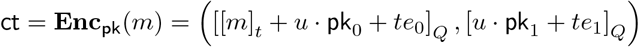

where *u* ← *χ*_ter_ and *e*_0_, *e*_1_ ← *χ*_err_. Due to the randomness in encryption, ciphertexts of the same plaintext *m* are always not identical, reflecting the IND-CPA security which avoids the search pattern leakage.

The relationship between a message *m* and its ciphertext (`ct`_0_, `ct`_1_) always satisfies

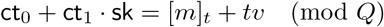

where *v* = *u* · *e* + *e*_1_ · `sk` + *e*_0_ is the noise term.

Only those knowing the secret key `sk` will be able to decrypt. Decrypting a ciphertext *ct* = (*ct*_0_, *ct*_1_) is to calculate

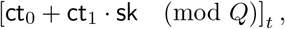

and it decrypts to the correct message *m* if the noise term *v* is lower than ⌊*Q/t*⌋.

#### Public Key-Switching

The key-switching technique is used to switch the secret key of a given ciphertext without decryption. Given a ciphertext ct that encrypts a message *m* under secret key `sk`, the goal is to obtain a new ciphertext that encrypts the same message *m* under another secret key `sk`^*^.

To compute this, the algorithm requires an additional ciphertext that encrypts the secret key `sk` under `sk`*, which is called the key-switching key 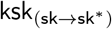. In the BGV scheme, this key is derived from *sk*^*^, but we propose an alternative derivation using `pk*`, the corresponding public key of `sk*`, as follows:

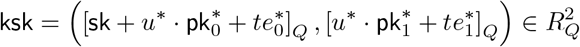

where 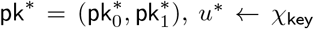 and 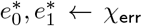. In other words, ksk is just **Enc**_pk_* (sk), i.e. the encryption of sk under pk^*^.

With ksk and (`ct`_0_, `ct`_1_) which decrypt to *m* under sk, we can construct 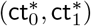 which also decrypt *m* but under `sk*`. Precisely, we construct

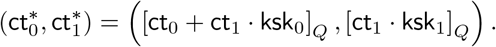

Therefore,

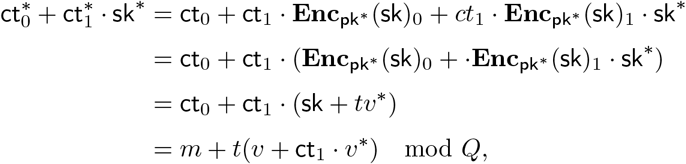

where 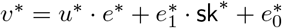 verifying 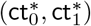 which also decrypts *m* but under `sk*`.

#### Reducing noise during public key-switching

The public key-switching above has noise component *ct*_1_ · *v*^*^, whose size can be further reduced by two methods: coefficient digit decomposition [18], and a temporary enlargement of ciphertext modulus [32]. We combine the two methods in an approach similar to the Section 4.3 of [37], where a ciphertext of modulus 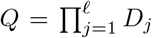 is decomposed into *𝓁* coprime and odd digits and the expansion factor *P* is also odd and coprime to *Q*.

As such, the key-switching key ksk_(sk→sk*)_ is no longer a ciphertext with two components, but a matrix of dimension 2 *× 𝓁* whose *j*-th column is 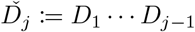, where *Ď*_*j*_ := *D*_1_ … *D*_*j*−1_ is the product of digits up to but not including *D*_*j*_. While [37] uses `sk*` to generate the key-switching matrix, our approach only requires `pk*` in the new key system.

#### Homomorphic operations

Adding two ciphertexts `ct` = (ct_0_, ct_1_) and 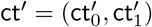 that encrypt *m* and *m*^*′*^ with a same key sk respectively gives

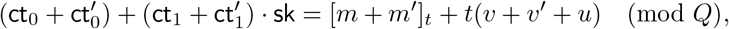

so the ciphertext 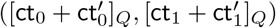 is an encryption of *m* + *m*^*′*^ (mod *t*) under almost additive noise.

Multiplication of two ciphertexts is more complex, which involves taking the tensor product of two ciphertexts as polynomial vectors to obtain 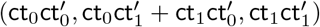. One can check that

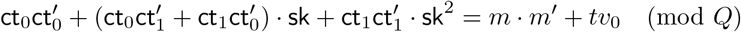

where *v*_0_ = *m* · *v*^*′*^ + *m*^*′*^ · *v* + *v* · *v*^*′*^.

Here, an additional step is needed for the term with `sk`^2^. If we consider 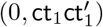 as a ciphertext with secret key `sk`^2^, then performing the key switching procedure with the key 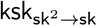 gives a ciphertext 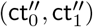. The noise control in this step is similar to the previous section. The final output of the homomorphic multiplication is 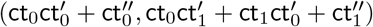.

#### SIMD Batching

The BGV scheme supports homomorphic operations on multiple plaintext slots simultaneously. This follows from the fact that the cyclotomic polynomial *Φ*_*m*_ (*X*) of degree *n* splits modulo *t* into *𝓁* irreducible factors of same degree 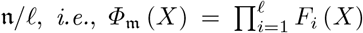. Leveraging the Chinese Reminder Theorem (CRT), the following ring isomorphism is established.

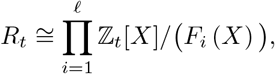

which enables the encoding of *𝓁* messages 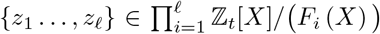 into a single plaintext in *R*_*t*_. Typically, each of the *𝓁* messages is called a *slot*, and altogether they are regarded as a length-*l* vector. Since computations over a ciphertext are performed on all packed values, BGV provides efficient computations in an amortized manner.

#### The Homomorphic rotation

BGV supports an additional operation called *rotation*, which permutes plaintext slots circularly. Specifically, let ct be an encryption of a plaintext vector **z** = (*z*_1_, *z*_2_, …, *z*_*𝓁*_). Performing a (right) rotation on ct by *v* results in a new ciphertext ct_rot_ encrypting the plaintext vector **z**_rot_ = (*z*_*v*+1_, *z*_*v*+2_, …, *z*_*𝓁*_, *z*_1_, …, *z*_*v*_) under the same secret key.

The homomorphic rotation consists of two key components: automorphism and key switching. We refer interested readers to Section 3 of [37] for the general hypercube structures of rotations in BGV.

### 7.11 Polynomial Evaluation using the Paterson-Stockmeyer method

Polynomial evaluations require numerous binary operations including additions, scalar multiplications (where one operand is a constant), and non-scalar multiplications. The Paterson-Stockmeyer method [54] uses fewer non-scalar multiplications, such that a degree-*d* polynomial is evaluated using 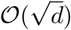 non-scalar multiplications.

In the homomorphic evaluation of polynomials, non-scalar multiplications are translated to ciphertext-ciphertext multiplications, whose evaluation costs are much more expensive than the other two. According to the benchmark in [30], it is around 160*×* and 15*×* the cost of homomorphic evaluations of additions and scalar multiplications, respectively. Therefore, the Paterson-Stockmeyer method has been widely used for polynomial evaluations in homomorphic encryption. [21,24,25,40].

Below we follow [25] to sketch the evaluation of a degree-*d* polynomial 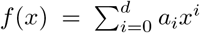 using the Paterson-Stockmeyer method. Assume there exist integers *L* and *H* such that *d* = *LH* −1 and 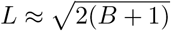. Then the polynomial can be rewritten into

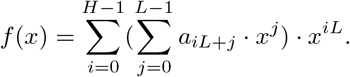

Therefore, the “low powers” {*x, x*^2^, …, *x*^*L*−1^} can be computed with *L* − 2 non-scalar multiplications, whose linear combinations give the inner sum. The “high powers” {*x*^*L*^, *x*^2*L*^, …, *x*^(*H*−1)*L*^} are then computed with *H* − 1 non-scalar multiplications, whose subsequent products with the inner sum require another *H* − 1 non-scalar multiplications. In total, the procedure requires

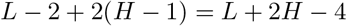

non-scalar multiplications, which is minimal and achieves 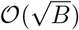 when 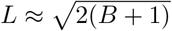.

### 7.12 Comparison of SQUiD’s MAF Calculation with Existing Methods

In the MAF protocol by Kim and Lauter [45], the MAF for SNP *j* is computed as follows:

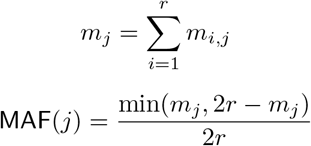

where *r* represents the number of patients in the database, and *m*_*i,j*_ denotes the genotype of patient *i* at SNP *j*

In SQUiD, a cohort is initially created before the MAF computation. The membership of the cohort is defined by a predicate vector *p*, which is 1 for patients in the cohort and 0 otherwise. The key distinction in the MAF computation between SQUiD and [45] lies in the need for multiplication of each *m*_*i,j*_ term within the summation by the corresponding predicate, ensuring the inclusion of only those patients who are part of the cohort. Additionally, the total number of patients *r* in the denominator is a constant in [45], but in SQUID it varies for different cohort sizes and needs to be computed as the sum of predicates. All other parts of the MAF computation are the same in both SQUiD and [45]. That is, a SQUiD MAF query without a filter has the same computation and result as [45]. Furthermore, in both SQUiD and [45], the division and minimum operations are executed in plaintext. In summary, the total runtime and accuracy of the SQUiD MAF calculation and that of [45] are expected to be exactly the same.

### 7.13 Continuous phenotype values

SQUiD supports continuous phenotype values such as weight, blood pressure, heart rate, etc. However, as SQUiD exclusively processes integer data inputs, these values need to be discretized into integers by scaling the values. This discretization occurs during the data encryption by the data owner before transmitting it to the cloud. Once in the cloud, SQUiD effectively filters these now integer values for counting and MAF queries using range filters. Range filters are set by an upper and lower bound and are computed using the same comparisons thresholds from the similarity query.

We benchmarked the range query performance in Supplementary Figure 8. We found that it took approximately 25 minutes to compute a count query with a range filter on 16,384 patients and 28 minutes to compute a MAF query with the same parameters.”

**Supplementary Figure 1:**
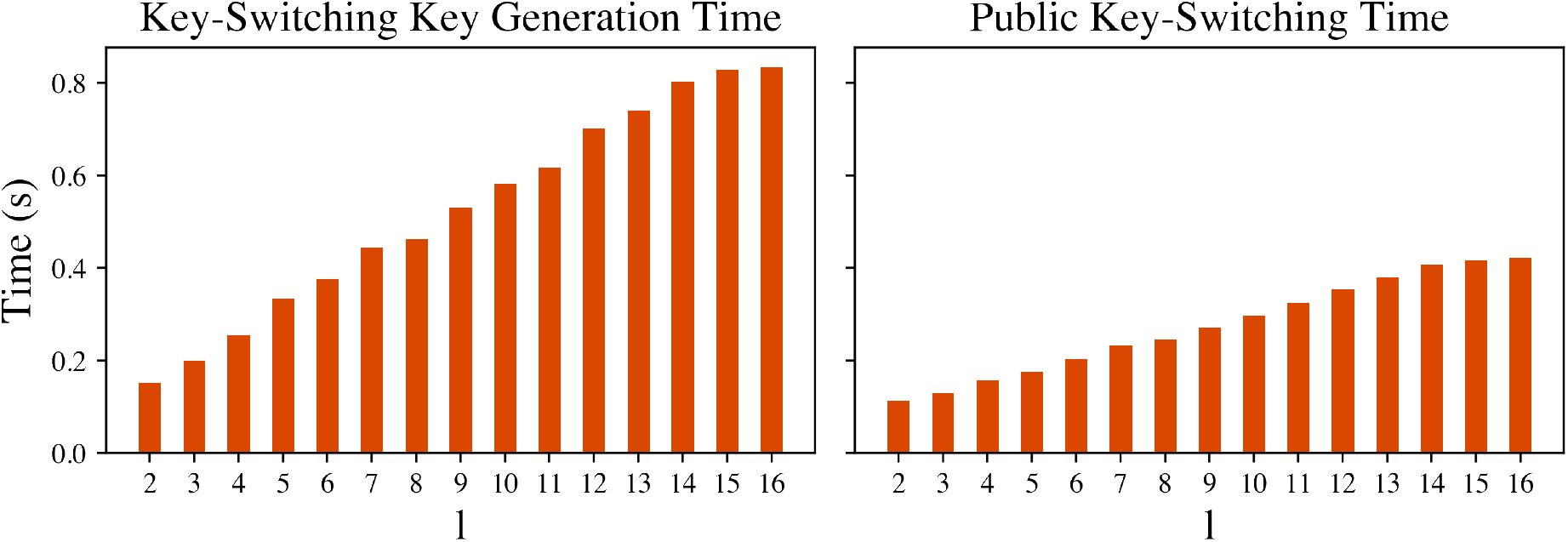
The runtime for key-switching key generation and key-switching by the number of digits used in the decomposition (l). Digit decomposition is a technique to reduce the error growth in ciphertexts. The more digits we decompose the ciphertext and key-switching key to, the more secure our system is and the less error growth the ciphertext experiences[38]. Overall, it 200 ms to generate a key-switching key when the digits are decomposed into three parts.

**Supplementary Figure 2:**
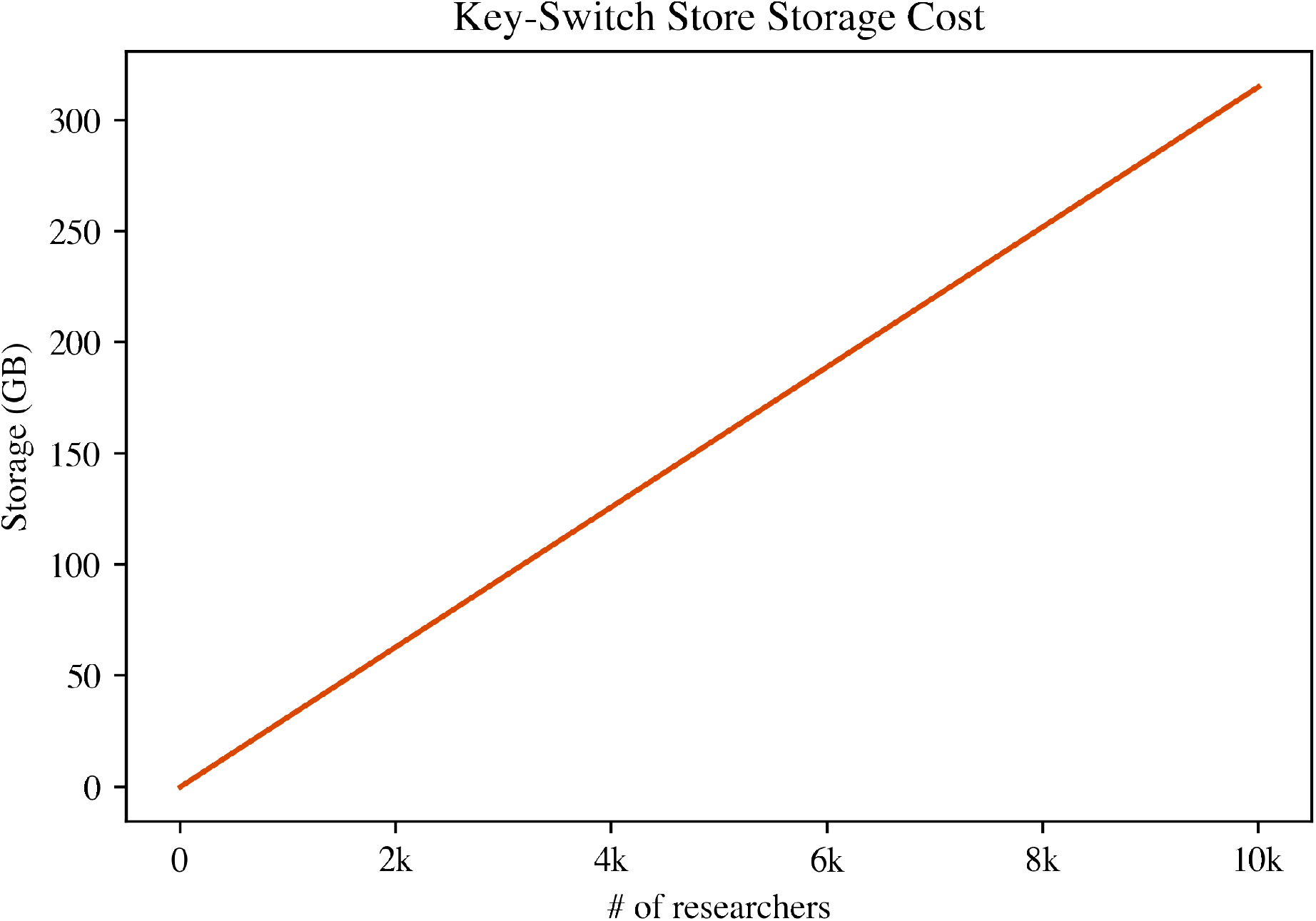
The additional storage space for the public key-switching key store (with l = 3) by the number of authorized researchers in the store. Since each key-switching key is a single ciphertext, the storage required remains minimal at approximately 31 gigabytes (GB) for 1000 researchers.

**Supplementary Figure 3:**
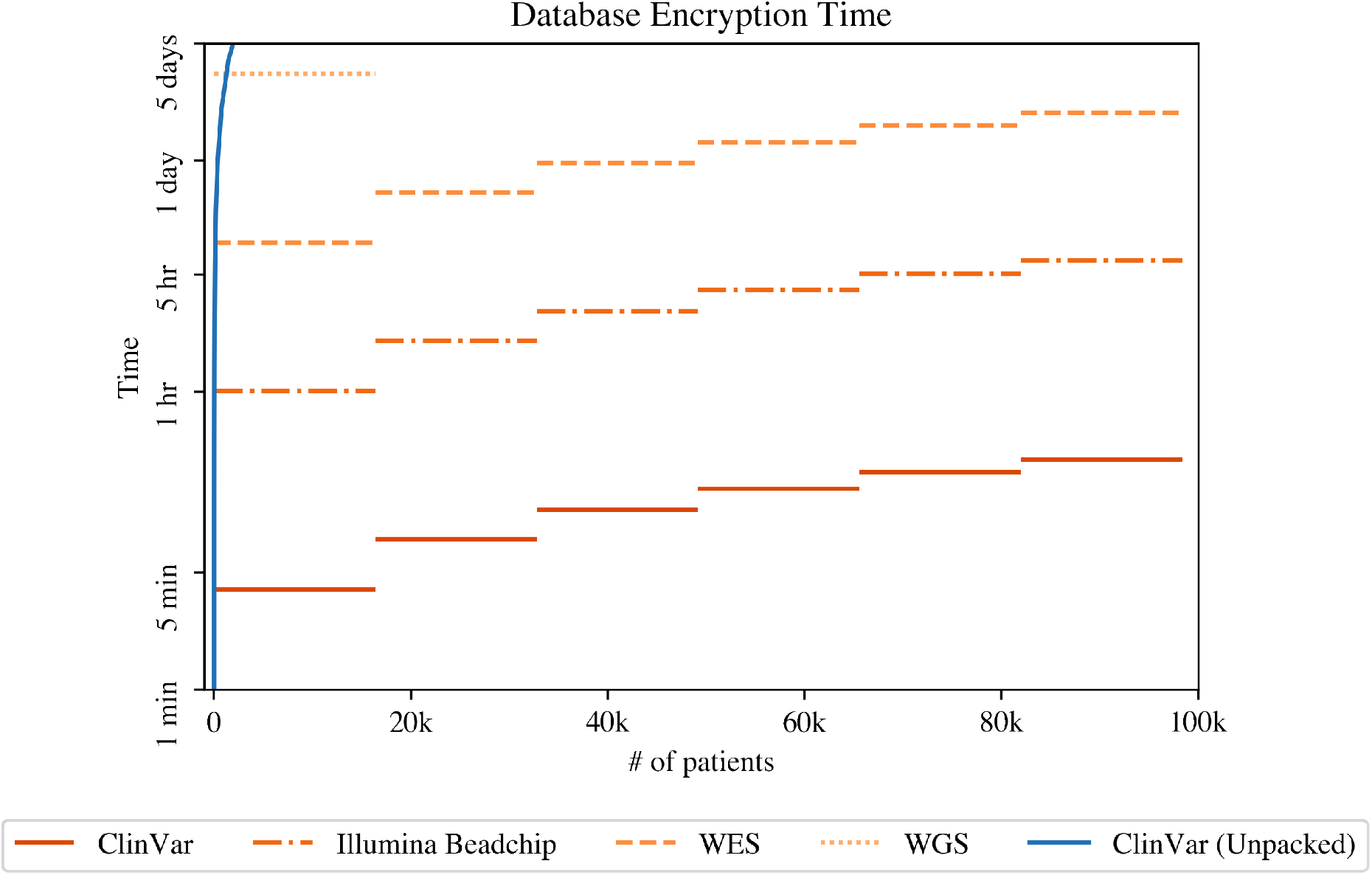
The time to encryption databases with various SNP sets by the number of patients in the database. We measured the setup time by encrypting the various databases using 35 threads running simultaneously. We compared the setup of SQUiD for the ClinVar SNP set to the setup time of an unpacked HE solution (Blue) for the ClinVar SNP set.

**Supplementary Figure 4:**
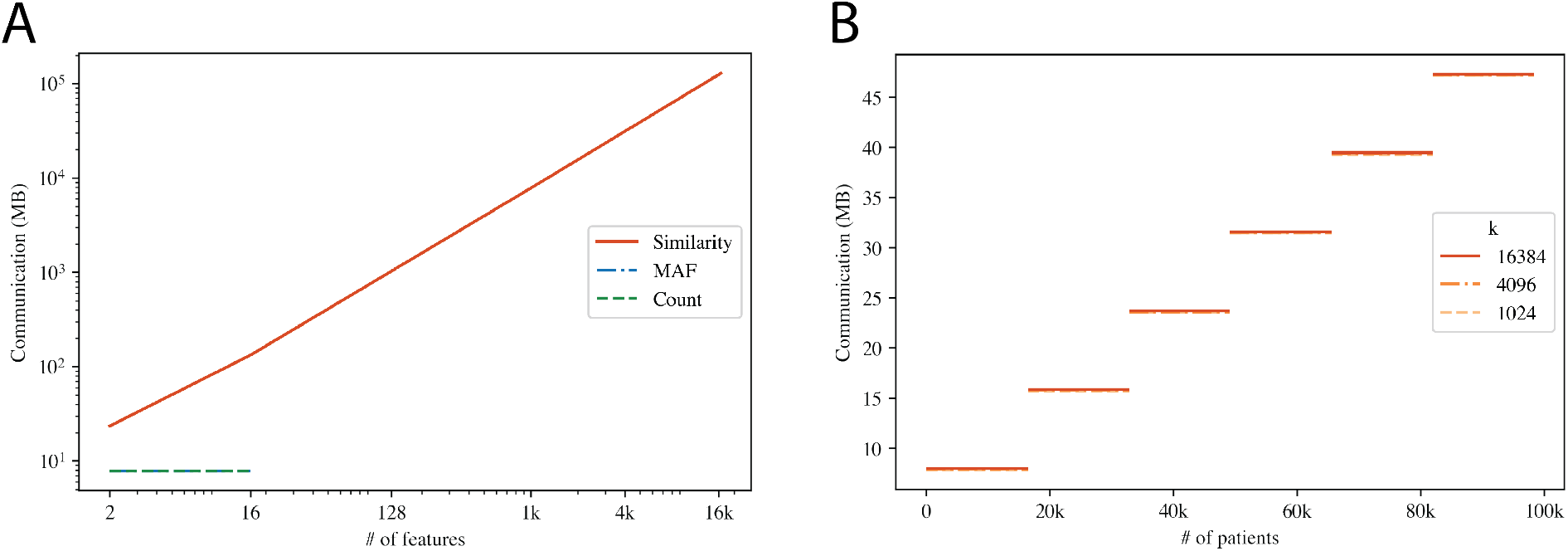
The communication cost for all queries by the number of features in the query or the number of patients in the database. We benchmark our communication cost by measuring the end-to-end communication of a single query. Each query needs one communication round thus the total communication is the cost of receiving a query from a client and sending back the computation result. We separated the communication performance into two plots because the scaling for our queries depend on different factors. (A) We show the communication costs of the count, MAF, and similarity queries by the number of features (filters for the count and MAF query, and SNPs for the similarity query). (B) We show the communication of the PRS query by the number of patients in the database.

**Supplementary Figure 5:**
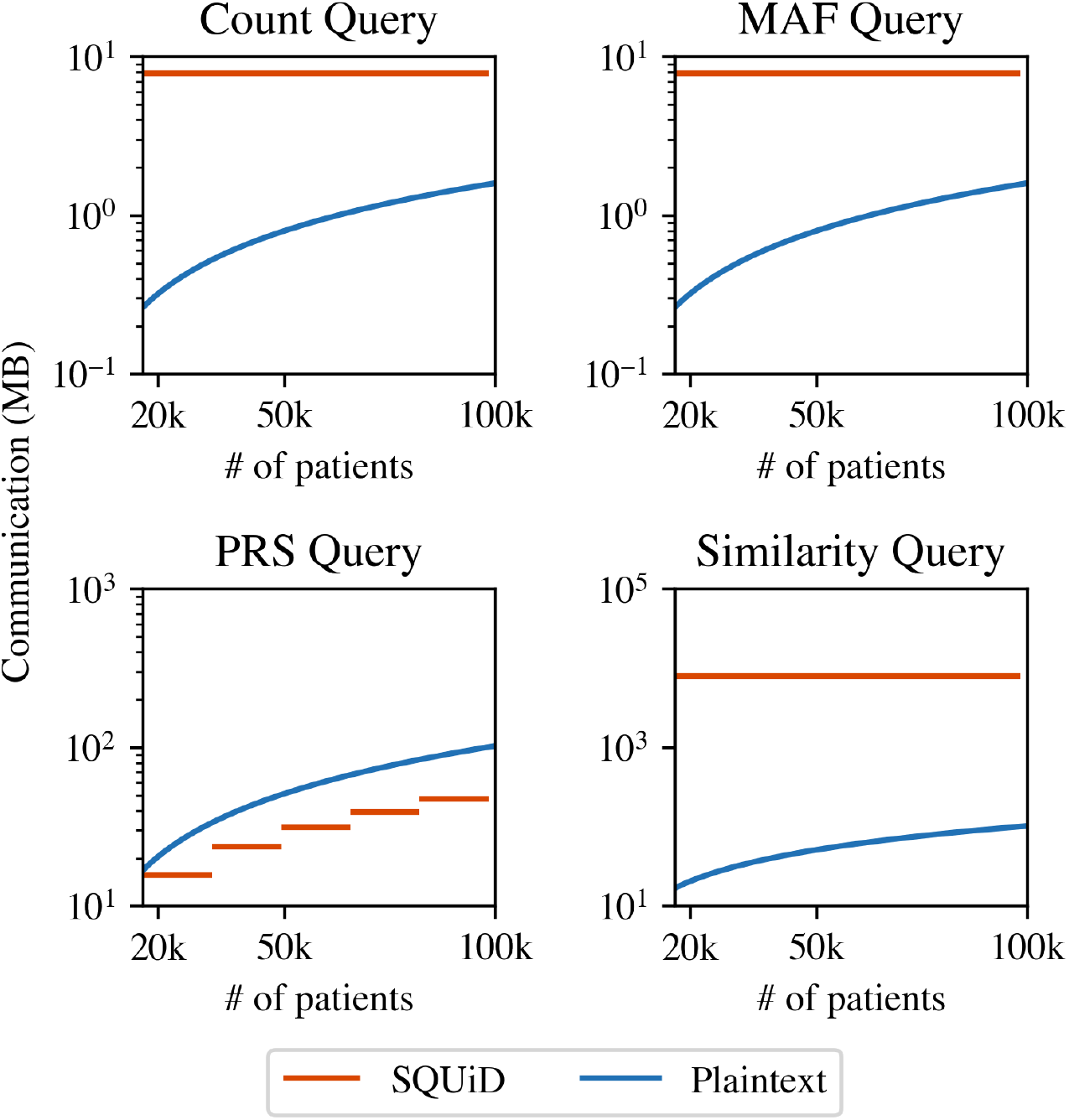
The communication cost for all queries and a plaintext solutions by the number of patients in the database. The plaintext database keeps the data encrypted at rest, packing 16 one byte SNPs into a single encrypted 128-bit AES block. When the database receives a query, it sends the encrypted data necessary to compute the query. Thus, the client has to decrypt and compute the query themselves. We benchmarked for all four types of queries with 16 filters for the count and MAF queries, 1024 effect sizes for the PRS query, and the 1,024 SNPs for the similarity query. Compared to SQUiD, the plaintext communication always scales linearly with the number of patients while this is only true for SQUiD for PRS queries (since a PRS score for each patient needs to be returned).

**Supplementary Figure 6:**
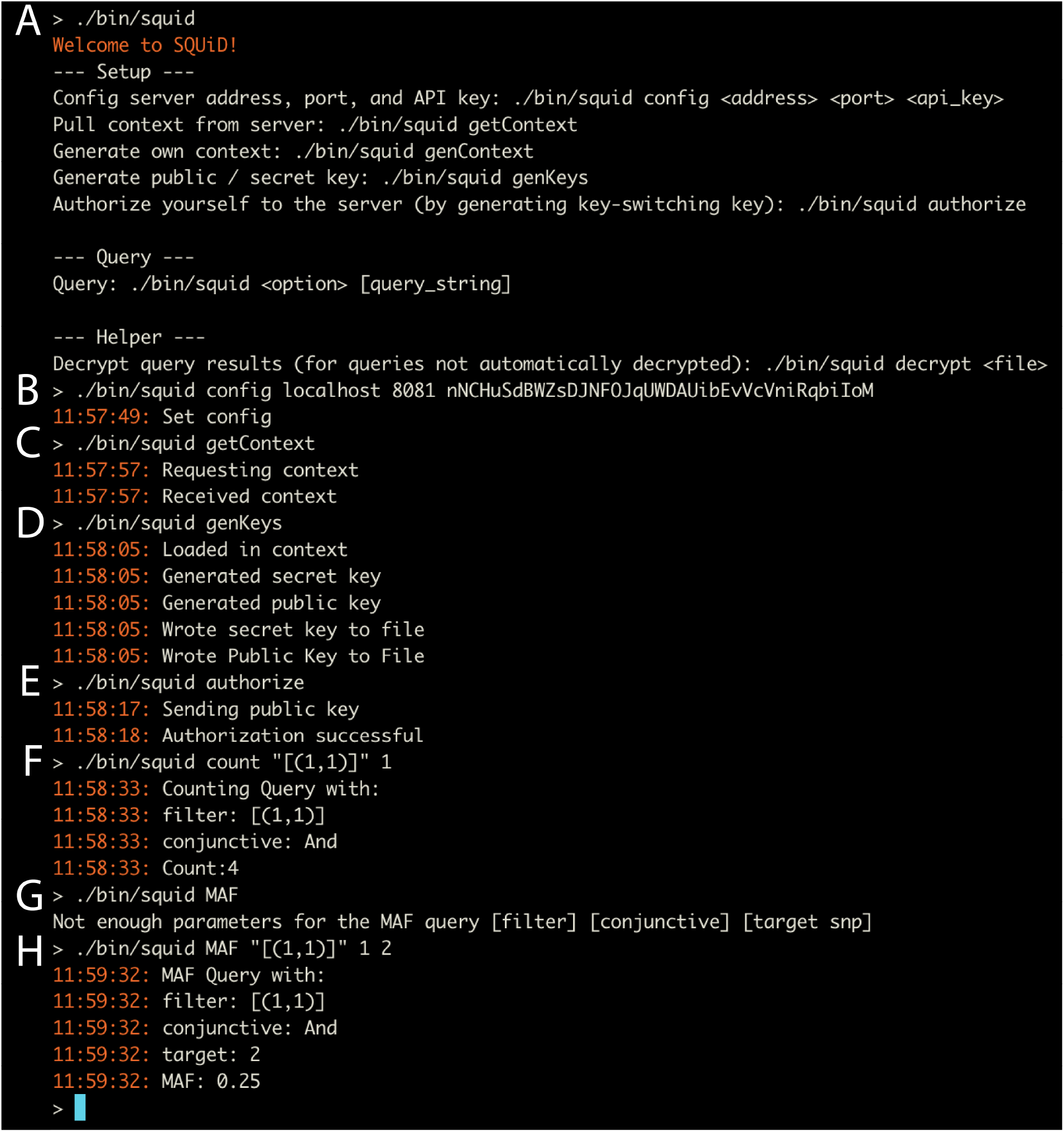
Screenshot of SQUiD command line interface (CLI). **(A)** Welcome help message about showing core SQUiD functionalities. **(B)** setConfig call sets the address, port, and API key used to communicate with the server for all successive calls. **(C)** getContext call that pulls the context from the server to ensure the encryption schemes are synced locally and on the server. **(DC** genKeys call creates a public and private key for the user. **(E)** authorize call sends the public key to the server for authorization. The server generates a key-switching key which will be used to re-encrypt all query results sent back to the user to be encrypted under the user’s public key. **(F)** count query call which counts the number of patients in the database with a first SNP that has value 1. **(G)** failed query call that shows what parameters should be supplied for a correct query call. **(H)** MAF query call that has the correct parameters.

**Supplementary Figure 7:**
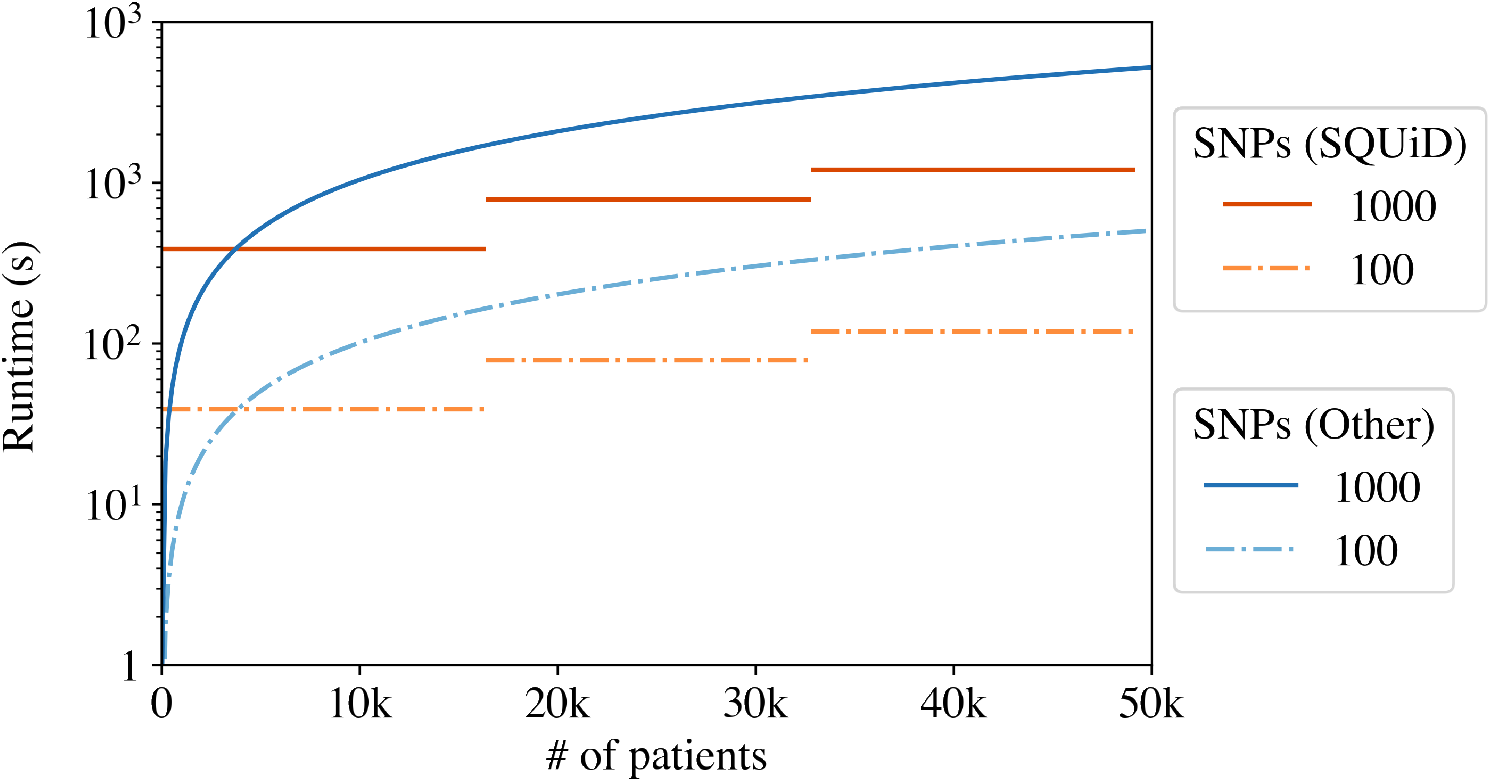
The runtime of the L2 similarity score computation for SQUiD and [59] (Other) for 100 SNPs and 1,000 SNPs. For score computation with fewer than 4,000 patients, [59] exhibits faster performance for both 100 and 1,000 SNPs. However, as the patient dataset scales up, SQUiD consistently outperforms [59], demonstrating its superior efficiency in handling larger datasets.

**Supplementary Figure 8:**
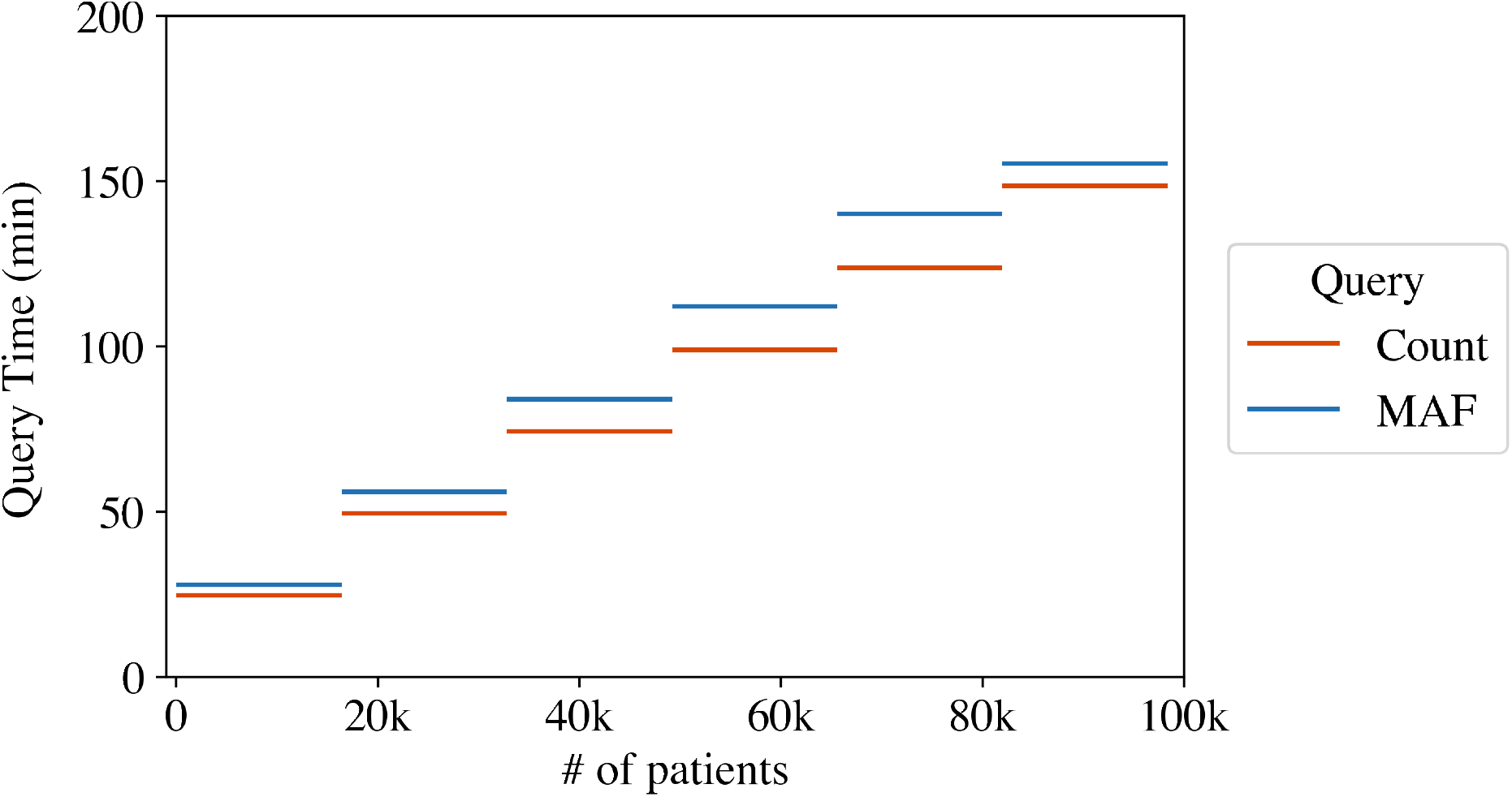
The query time for the count and MAF query with a range filter by the number of patients in the database. Each query only had one range filter.

